# Pangenome Graph Construction from Genome Alignment with Minigraph-Cactus

**DOI:** 10.1101/2022.10.06.511217

**Authors:** Glenn Hickey, Jean Monlong, Jana Ebler, Adam Novak, Jordan M. Eizenga, Yan Gao, Human Pangenome Reference Consortium, Tobias Marschall, Heng Li, Benedict Paten

**Affiliations:** UC Santa Cruz Genomics Institute, University of California, Santa Cruz, 1156 High St, Santa Cruz, CA, USA; Institute for Medical Biometry and Bioinformatics, Medical Faculty, Heinrich Heine University Düsseldorf, Düsseldorf, Germany; Center for Digital Medicine, Heinrich Heine University Düsseldorf, 40225 Düsseldorf, Germany; Center for Computational and Genomic Medicine, The Children’s Hospital of Philadelphia, Philadelphia, PA 19104, USA.; Department of Data Sciences, Dana-Farber Cancer Institute, Boston, MA 02215, USA; Department of Biomedical Informatics, Harvard Medical School, Boston, MA 02215, USA

## Abstract

Reference genomes provide mapping targets and coordinate systems but introduce biases when samples under study diverge sufficiently from them. Pangenome references seek to address this by storing a representative set of diverse haplotypes and their alignment, usually as a graph. Alternate alleles determined by variant callers can be used to construct pangenome graphs, but thanks to advances in long-read sequencing, high-quality phased assemblies are becoming widely available. Constructing a pangenome graph directly from assemblies, as opposed to variant calls, leverages the graph’s ability to consistently represent variation at different scales and reduces biases introduced by reference-based variant calls. Pangenome construction in this way is equivalent to multiple genome alignment. Here we present the Minigraph-Cactus pangenome pipeline, a method to create pangenomes directly from whole-genome alignments, and demonstrate its ability to scale to 90 human haplotypes from the Human Pangenome Reference Consortium (HPRC). This tool was designed to build graphs containing all forms of genetic variation while still being practical for use with current mapping and genotyping tools. We show that this graph is useful both for studying variation within the input haplotypes, but also as a basis for achieving state of the art performance in short and long read mapping, small variant calling and structural variant genotyping. We further measure the effect of the quality and completeness of reference genomes used for analysis within the pangenomes, and show that using the CHM13 reference from the Telomere-to-Telomere Consortium improves the accuracy of our methods, even after projecting back to GRCh38. We also demonstrate that our method can apply to nonhuman data by showing improved mapping and variant detection sensitivity with a *Drosophila melanogaster* pangenome.

## Introduction

The term pangenome has historically referred to the set of genes present across a population or species. The patterns of presence and absence of genes from the pangenome in individual samples, typically prokaryotes, provided a rich context for better understanding the genes and populations in question ^1^. Eukaryotic genomes can likewise be combined into pangenomes, which can be expressed in terms of genomic content rather than genes. Eukaryotic pangenomics is growing in popularity, due in part to its potential to reduce reference bias ^2^.

A pangenome can be represented as a set of variants against a reference ^3^, but technological advances in long-read sequencing are now making it possible to produce high-quality *de novo* genome assemblies of samples under study, allowing for variation to be studied within its full genomic context ^4^. Two themes that have emerged from this work are that 1) relying on a single reference genome can be a source of bias, especially for short-read sequencing projects, and 2) representation of structural variation is a challenging problem in its own right. Pangenomes and the software toolkits that work with them aim to address these issues.

Sequence-resolved pangenomes are typically represented using graph models. There are two main classes of graph representation: sequence graphs and de-Bruijn graphs, and several different methods have been published for each type. This is an area of active research; different methods perform better for different applications, and there is as yet no clear best practice. However, sequence graphs have generally proved more amenable for read mapping ^3, 5, 6^, and they will be the focus of this work. In a sequence graph, each node corresponds to a DNA sequence (**Figure 1A**) or its reverse complement depending on the direction in which it is traversed. Sample haplotypes are stored as paths, and edges are bidirected to encode strandedness (i.e. if an edge is incident to the forward or reverse complement sequence of a node). Sites of variation appear as bubbles, or snarls, which are defined by characteristic subgraphs ^7^. Two snarls are indicated in the example graph in **Figure 1A,** the left and right representing a two-base substitution and 19-base deletion, respectively.

**Figure 1:**
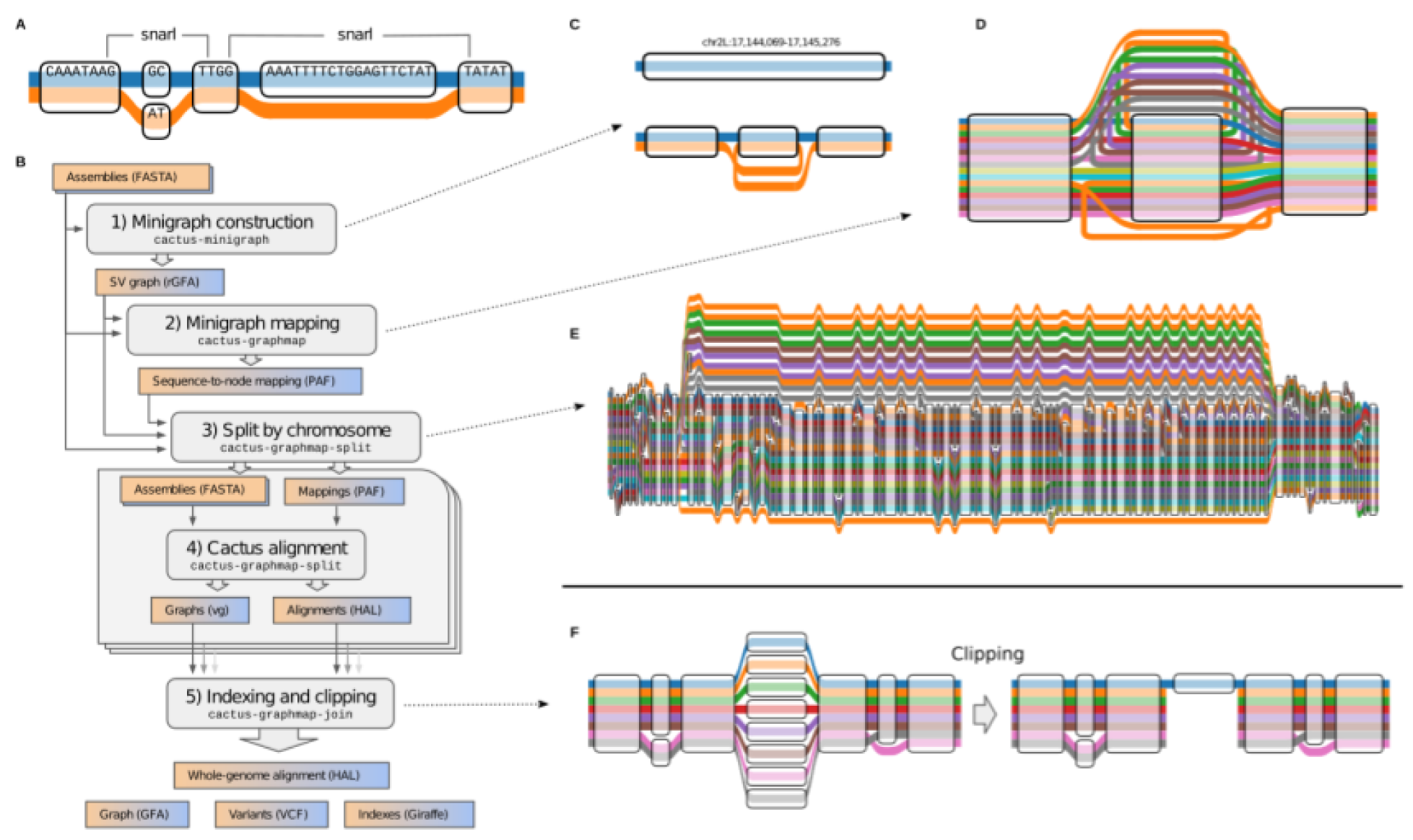
Minigraph-Cactus Pangenome Construction. **A)** “Tube Map’’ view of a sequence graph shows two haplotypes as paths through the graph. The two snarls (variation sites defined by graph topology, aka bubbles) are highlighted. **B)** The five steps, and associated tools, of the Minigraph-Cactus pipeline which takes as input genome assemblies in FASTA format and outputs a pangenome graph, genome alignment, VCF and indexes required for mapping with vg Giraffe. Illustrating the steps in the pipeline by example: **C)** SV graph construction using minigraph (as wrapped by cactus-minigraph) begins with a linear reference and adds SVs, in this case a single 1204bp inversion (at ch2L:17,144,069 in the *D. melanogaster* pangenome). **D)** The input haplotypes are mapped back to the graph with minigraph, in this example six of which contain the inversion allele from C. **E)** The minigraph mappings are combined into a base-resolution graph using Cactus, augmenting the larger SVs with smaller variants - in this case, adding smaller variants within the inversion. **F)** An unaligned centromere is clipped out of a graph, leaving only the reference (blue) allele in that region. The other alleles are each broken into two separate subpaths but are otherwise unaffected outside the clipped region.

Phased Variant Call Format (VCF) files can be thought of as sequence graphs. The vg toolkit makes this perspective explicit by supporting graph construction from VCF ^3^. Using such graphs for mapping and variant calling reduces reference bias and improves accuracy over GRCh38 ^3, 6^. These graphs can also be used to accurately genotype structural variants (SVs) ^5^, but they are still limited to reference-based variant calls. For example, there is no satisfactory way in VCF 4.3 to directly represent variation nested within a large insertion. Now that they are becoming widely available^8^, high-quality assemblies can instead be used to directly construct a pangenome graph without the need to go through variant calls. This is equivalent to finding a whole genome multiple alignment, which is known to be an extremely computationally challenging problem ^9^.

As such, multiple alignment algorithms must use heuristics for scaling with respect to both the number of input sequences and their combined length. Typically, the former is accomplished by decomposing the multiple alignment of N genomes into smaller subalignments that can be composed together, and the latter by seed-and-extend heuristics ^10^.

MultiZ ^11^ was among the first methods able to align dozens of vertebrate genomes and is still used by the UCSC Genome Browser. It begins with a set of pairwise alignments of the input genomes to a given reference assembly, then uses progressive decomposition to merge the alignments according to their phylogenetic relationships. The pairwise alignments themselves are created with LASTZ, which uses a gapped seeding approach to find anchors, which are then chained and extended with dynamic programming ^12^. Progressive Cactus is a more recent and scalable tool for large vertebrate scale multiple alignments ^13^. It also uses LASTZ, or the GPU-accelerated successor SegAlign ^14^, to perform pairwise alignments. However, it does so by progressively reconstructing ancestral sequences using a phylogenetic guide tree. This eliminates the need for a global reference assembly, making Progressive Cactus reference-independent. At each step, the LASTZ alignments are used as anchors to construct a cactus graph ^15^, which in turn is used to filter and then refine the alignment.

Progressive Cactus was shown to be robust to small errors in the guide tree, but, like any progressive alignment approach, it still relies upon an accurate phylogenetic tree. Due to recombination, a single tree cannot reasonably represent the ancestry of any intraspecies genome set that one might want to use to construct a pangenome. Minigraph ^16^ is a newer tool that uses an iterative sequence-to-graph mapping approach, similar to Partial Order Alignment (POA) ^17^, to construct a pangenome graph from a set of input genomes. It uses a generalization of minimap2’s minimizer-based seeding and chaining strategy ^18^, and is similarly fast so long as the input genomes are relatively similar. While minigraph can perform base-level alignment since version 0.17, it only includes SVs (≥ 50bp by default) during graph construction. Excluding small variation prevents input genomes from being losslessly embedded as paths in the graph, as well as the joint consideration of all types of variants with a single model.

We now present Minigraph-Cactus, a new pangenomics pipeline that combines Minigraph’s fast assembly-to-graph mapping with a novel version of Cactus’s base aligner, alongside several key improvements in vg ^3, 6^), in order to produce base-level pangenome graphs at the scale of dozens to hundreds of vertebrate haplotypes. In addition to representing variation consistently at all resolutions, we show that these graphs can be used to improve upon the state of the art for short and long-read mapping, variant calling, and SV genotyping.

## The Minigraph-Cactus Pangenome Pipeline

The Minigraph-Cactus Pangenome pipeline has been added to the Cactus software suite. Like Progressive Cactus ^13^, it is implemented using Toil ^19^, which allows it to be run either locally or via distributed computation on clusters, including those provisioned in the cloud. The pipeline consists of five steps as shown in **Figure 1B**, which are used to generate a graph in both GFA and VCF format, as well as indexes required to map reads using vg giraffe ^6^.

## Minigraph SV graph construction

The pipeline begins with the construction of an initial SV-only graph using minigraph as described in ^16^. By default, only variants affecting 50bp of sequence or more are included. This is an iterative procedure that closely resembles partial order alignment (POA): a “reference” assembly is chosen as an initial backbone, and then augmented with variation from the remaining assemblies in turn. **Figure 1C** shows an example of an inversion being augmented into a reference chromosome. Minigraph does not collapse duplications: If two copies of a gene are present in the graph after adding *i* genomes, but there are three copies in the *i+1*th genome, then an additional copy will be added to the graph. This is a key difference between minigraph and other approaches (including Progressive Cactus) that would tend to collapse all copies of the gene into a single sequence in the absence of outgroup information to determine the ancestral state. By keeping different gene copies separate, minigraph trades greater graph size for reduced path complexity (fewer cycles).

## Minigraph contig mapping

Minigraph generalizes the minimizer-based seeding and chaining concepts from minimap2 ^18^ for use on sequence graphs. For this current work we generalized it to produce base-level alignments between contigs and graphs (but not base-level *graphs*). In this step of the pipeline each assembly, including the reference, is mapped back to the SV graph independently (**Figure 1D)**. The results are concatenated into a single Graphical Alignment Format (GAF) file, which is then filtered to remove spurious alignments (See Methods Section for details). By re-aligning each assembly to the same graph in this step as opposed to re-using the iterative mappings created during construction, we mitigate an issue in the latter where orthologous sequences can be aligned to inconsistent locations when mapped to different versions of the graph.

## Splitting by chromosome

Minigraph does not introduce interchromosomal events during graph construction, so every node in the SV graph is connected to exactly one chromosome (or contig) from the reference assembly. This information is used to split the *mappings* obtained in the previous step into chromosomes. If a contig maps to nodes from multiple chromosomes, it is assigned to the chromosome to which the most of its bases align. Thresholds (detailed in the Methods Section) are used to filter out contigs that cannot be confidently assigned to any reference chromosome. Such contigs will be excluded from the constructed graph. Graph construction proceeds on each reference chromosome independently, which serves to increase parallelism and reduce peak memory usage (per job). These computational advantages are required to construct a 90- sample human pangenome graph on current hardware, but smaller datasets could be run all at once if desired, avoiding this step entirely.

## Cactus base alignment

At its core, Cactus is a procedure for combining a set of pairwise alignments into a multiple alignment ^13, 20^: It begins by “pinching” exactly matching aligned bases together in the pairwise alignments to form an initial sequence graph (**Figure 1A**). This sequence graph is then transformed into a Cactus graph (**Supplementary Figure 1A-C**), whose cycles represent the “chains” of alignment within the sequence graph ^15^. The topology of the Cactus graph is first used to remove candidate spurious or incomplete alignments corresponding to short, high-degree alignment chains. Interstitial unaligned sequences that share common anchors at their ends are then aligned together. This process as a whole remains unchanged at a conceptual level when using Cactus to construct pangenome alignments, but substantial changes to each step were required by the increase in the number of input genomes: Cactus does not typically align more than four genomes (two ingroups and two outgroups) at a time when computing progressive alignments, so scaling to 90 HPRC samples (and beyond) required the underlying graph structures to be rewritten to use less memory, as well as completely replacing the algorithm for interstitial sequence alignment. Briefly, the previous all-pairs approach, which scales quadratically with the number of genomes, was replaced with a Partial Order Alignment (POA) approach that scales linearly (See Methods for details).

Cactus natively outputs genome alignments in Hierarchical Alignment (HAL) format ^21^. HAL files can be used to create assembly hubs on the UCSC genome browser, or to map annotations between genomes ^22^, but they are not suitable for most pangenome graph applications, which expect GFA or VG. We therefore created a new tool, hal2vg, to convert HAL alignments into VG format. (see Methods for more details). These graphs contain the underlying structural variation from the SV graph constructed by minigraph along with smaller variants, and the input haplotypes are represented as paths (**Figure 1E**).

## Indexing and clipping

The final step of the pipeline combines the chromosome level results and performs some post-processing. This includes reassigning node ids so that they are globally unique across different chromosome graphs, and collapsing redundant sequence where possible using gaffix ^23^(**Supplementary Figure 1D**). Nodes are also replaced with their reverse complement as necessary to ensure that reference paths only ever visit them in the forward orientation. The original SV graph produced by minigraph remains embedded in the results at this stage, with each minigraph node being represented by a separate embedded path.

Minigraph-Cactus (in common with all MSA tools we know of^24^) cannot presently satisfactorily align highly repetitive sequences like satellite arrays, centromeres and telomeres because they lack sufficiently unique subsequences for minigraph to use as alignment seeds. As such, these regions will remain largely unaligned throughout the pipeline and will make the graph difficult to index and map to by introducing vast amounts of redundant sequence. We recommend clipping them out for most applications and provide the option to do so by removing paths with >N bases that do not align to the underlying SV graph constructed with minigraph (**Figure 1F**). In preliminary studies of mapping short reads and calling small variants (see below), we found that even more aggressively filtering the graph helps improve accuracy. For this reason, an optional allele-frequency filter is included to remove nodes of the graph present in fewer than N haplotypes and can be used when making indexes for vg giraffe.

In all, up to three graphs are produced while indexing:

1. Full graph: useful for storing complete sequences and performing liftover (translation between corresponding haplotypes); difficult to index and map to because of unaligned centromeres. These graphs are typically created only as intermediate results, and are not directly used in any of the results in this report.
2. Default graph: clip out all stretches of sequences >=10kb that do not align to the minigraph. The intuition is that large SVs not in minigraph are under-alignments of sequence not presently alignable and not true variants. The 10kb threshold is arbitrary but empirically was found to work well. This graph is ideal for studying variation and exporting to VCF, and can be effectively indexed for read mapping. These graphs are used in all results unless otherwise is explicitly stated.
3. Allele-frequency filtered graph: remove all nodes present in fewer than N haplotypes. This filter increases accuracy for short read mapping and variant calling, as shown in **Supplementary Figures 6** and **7**, respectively. These graphs are used for mapping with vg giraffe.

Graph 2) is a subgraph of graph 1), and graph 3) is a subgraph of graph 2). They are node-id compatible, in that any node shared between two of the graphs will have the same sequence and ID. Unless otherwise stated, all results below about the graphs themselves are referring to the default graphs, whereas all results pertaining to short read mapping and small variant calling were performed on the allele-frequency filtered graphs.

## Human Pangenome Reference Graphs

The Minigraph-Cactus pipeline was originally developed to construct a pangenome graph for the assemblies produced by the Human Pangenome Reference Consortium (HPRC). In its first year, this consortium released 47 diploid assemblies ^25^. For evaluation purposes, we held out three samples when generating the graph: HG002, HG005 and NA19240. The remaining 44 samples (88 haplotypes), and two reference genomes (GRCh38 and CHM13 [v1.1] ^26^ were used to construct the graph, with 90 haploid genomes total. Since the construction procedure is dependent on the reference chosen for the graph, we ran our pipeline twice independently on the same input assemblies, once using GRCh38 as the reference and once CHM13. The CHM13-based graph includes more difficult and highly variant regions, such as in the acrocentric short arm of chr21, that are not represented in the GRCh38-based graph. This makes it slightly bigger than the GRCh38-based graph, both in terms of total sequence and in terms of nodes and edges (**Supplementary Table 1**). The final pangenomes have roughly 200X more nodes and edges than the SV Graphs from Minigraph, showing the amount of small variation required in order to embed the haplotype paths. **Figure 2A** shows the amount of non-reference sequence as a function of how many haploid genomes contain it (the same plot for total sequence can be found in **Supplementary Figure 2**). The rise in the leftmost points (support=1) is due to private sequence, only present in one sample, and may also contain alignment artifacts which often manifest as under-alignments affecting a single sample. The plot clearly shows that the CHM13-based graph has less non-reference sequence present across the majority of samples, an apparent consequence of the improved completeness of CHM13 over GRCh38. The distribution of allele sizes within snarls (variant sites in the pangenome defined by graph topology; **Figure 2B**) highlights the amount of small variation added relative to Minigraph alone. The total time to create and index each HPRC pangenome graph was roughly 3 days (**Supplementary Table 4**).

**Figure 2:**
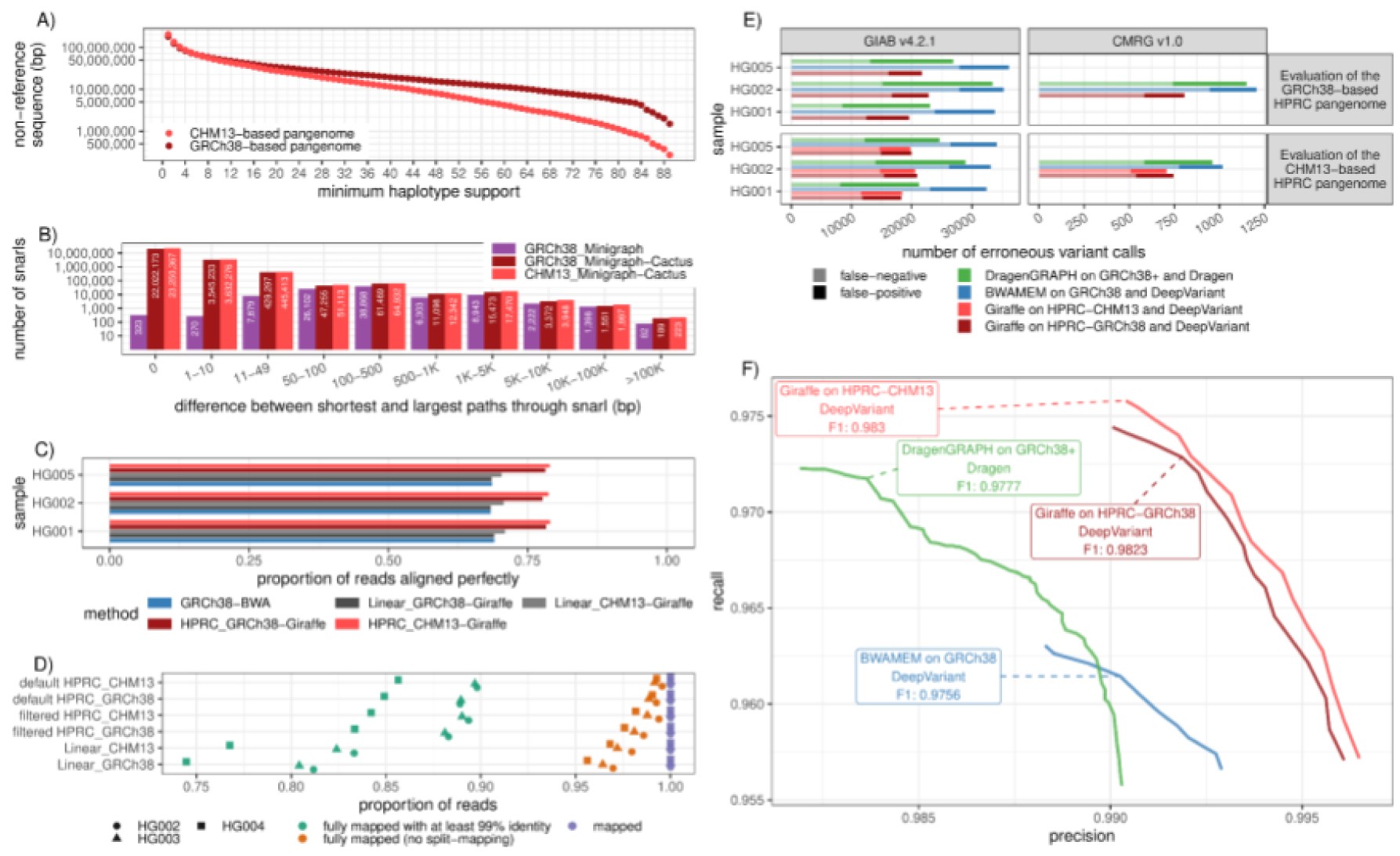
Evaluating GRCh38 and T2T-CHM13 based human pangenomes. **A)** The amount of non-reference sequence in the HPRC graphs by the minimum number of haplotypes it is contained in. **B)** Distribution of the size of the snarls (variation sites, aka bubbles) for the GRCh38-based Minigraph, GRCh38-based and CHM13-based Minigraph-Cactus pangenomes. Note that in the case of overlapping variants, snarls can be much larger than any single event that they contain. **C)** ∼30x Illumina short-reads for three GIAB samples were mapped using three approaches: BWA-MEM on GRCh38 (blue), vg Giraffe on the linear pangenomes with GRCh 38 or CHM13 (grey), vg Giraffe on the GRCh38-referenced or CHM13-referenced HPRC pangenome (red). **C)** Proportion of the reads aligning perfectly to the (pan-)genome for each sample (y-axis). **D)** Number of HiFi reads mapped to the linear, filtered, and default (unfiltered by allele frequency)pangenomes. For each sample and pangenome, three points show the number of mapped reads (purple square), reads mapped without being split (orange triangle), and reads fully mapped with at least 99% identity. **E-F)** Short variants were called with DeepVariant after projecting the reads to GCRh38 from the GRCh38-based pangenome (dark red), or the CHM13-based pangenome (light red). The results when aligning reads with BWA-MEM (blue) or using the Dragen pipeline (green) are also shown. **E)** The number of erroneous calls (false positive in dark, false-negative in pale) is shown on the x-axis across samples from the Genome in a Bottle (y-axis). Left: Genome in a Bottle v4.2.2 high confidence calls. Right: Challenging Medically Relevant Genes v1.0. When evaluating the CHM13-based pangenome (bottom panels), regions with false duplications or collapsed in GRCh38 were excluded. **F)** The graph shows the precision (x-axis) and recall (y-axis) for different approaches using the Challenging Medically Relevant Genes v1.0 truth set for the HG002 sample (bottom-right panel in **E)**). The curves are traced by increasing the minimum quality of the calls.

## Mapping to the HPRC Graphs

We benchmarked how well the pangenome graphs could be used as drop-in replacements for linear references in a state-of-the-art small variant (<50bp) discovery and genotyping pipeline. To do so, we used Illumina short reads (∼30x coverage) from three Genome in a Bottle (GIAB) samples, HG001, HG002, and HG005. All mapping experiments were performed on filtered HPRC graphs with a minimum allele frequency of 10%, meaning that nodes supported by fewer than 9 haplotypes were removed. This threshold was chosen to maximize variant calling sensitivity and mapping speed for the Giraffe-DeepVariant pipeline (**Supplementary Figures 8 and 9, respectively).** We found that reads aligned with higher identity when mapped to the pangenomes using Giraffe, compared to the traditional approach of mapping reads with BWA-MEM on GRCh38. We also mapped reads to the linear references with Giraffe and achieved similar results to using BWA. On average, 78.1% and 78.9% of reads aligned perfectly for the GRCh38-based and CHM13-based pangenomes, respectively, compared to 68.7% when using BWA-MEM on GRCh38 (**Figure 2C**). Similarly, reads mapped to the pangenomes had higher alignment scores (**Supplementary Figure 5**). Mapping to the pangenomes results in a slight drop in mapping confidence, from about 94.9% to 94.1% of reads with a mapping quality greater than 0 (**Supplementary Figure 6**) in those samples. This is expected as the pangenome contains more sequence than GRCh38, including complex regions and large duplications that are more fully represented, which naturally and correctly reduces mapping confidence for some reads. The same trend is observed when the pangenome is not filtered by frequency (**Supplementary Figure 6**). We also compared the alignment of long HiFi reads, mapped with GraphAligner^27^. Mapping to the pangenomes results in more long reads mapped fully (i.e. no split mapping) and with high identity (**Figure 2D**).

## Variant Calling with the HPRC Graphs

We used the short-read alignments to call variants with DeepVariant ^28^. To prepare them for DeepVariant, the graph alignments were projected onto GRCh38 using the vg toolkit. Note that, even though the CHM13-based graph did not use GRCh38 as the initial reference, the graph does contain GRCh38. Thus, the CHM13-based graph can also be used in this pipeline.

Both pangenomes constructed with Minigraph-Cactus outperform current top-performing methods (**Figure 2E-F**). We note that reads in regions that are falsely duplicated or collapsed in GRCh38 cannot be unambiguously projected from their corrected alleles in CHM13. For this reason, these regions were removed from the benchmark when evaluating the CHM13-based pangenome. Unsurprisingly, the CHM13-based pangenome offers the largest gains in variant calling in challenging regions like those assessed by the Challenging Medically Relevant Genes (CMRG) truth set (**Figure 2E**) ^29^. **Figure 1F** shows the precision and recall curves and the CHM13-based pangenome-based variant calls vs state of the art methods based on linear references for the CMRG benchmark. The CHM13-and GRCh38-based pangenomes have F1 scores 0.9830 and 0.9823, respectively, compared to 0.9777 and 0.9756 of Dragen and BWA-MEM DeepVariant, respectively. This gain in F1, though modest, still corresponds to hundreds of variants in these regions (**Figure 2E)**. The frequency-filtered pangenomes performed better than using the default pangenomes (**Supplementary Figure 7**). We also tested projecting and calling variants on CHM13. Although the benchmarking protocol is still preliminary for CHM13, we observed a clear improvement when using the pangenome compared to aligning the reads to CHM13 only (**Supplementary Figure 10**). Some specific regions, including the MHC region and segmental duplications, also have better variant calls on the CHM13-based graph (**Supplementary Figure 11**).

## Structural Variant Genotyping with the HPRC Graphs

PanGenie is a state-of-the art tool for genotyping human structural variation using short reads ^30^. It uses an HMM that combines information from known haplotypes in a pangenome (as represented by phased VCF) along with kmers from short reads in order to infer genotypes and, as such, does not require any read mapping. Minigraph-Cactus can output phased VCF representations of pangenome graphs that can be used as input to PanGenie (see Methods for more details). We evaluated this process by genotyping a cohort of 368 samples from the 1000 Genomes Project^31^ (1KG) comprising 20 trios randomly selected from each of the five superpopulations, along with the samples present in the graphs. We repeated this process independently on three different graphs: the GRCh38-based and CHM13-based HPRC pangenomes, as well the v2.0 PanGenie lenient variant set produced by the Human Structural Variation Consortium (HGSVC) ^32^. This latter graph was made by constructing reference-based variant calls for each sample, then merging similar variants together into single consensus variants, exactly the process that our pipeline is designed to avoid. The number of variants in each graph is given in **Supplementary Table 3**.

In order to measure PanGenie’s accuracy on each graph, we performed a leave-one-out experiment on five samples from the graphs. For each selected sample, its genotypes and private variants were removed from the VCF, which was then re-genotyped with PanGenie using short reads from that sample. These genotypes were then compared back to those from the original graph, effectively measuring how closely the haplotypes from short-read genotyping correspond to the original, assembly-based haplotypes. Due to their disjoint sample sets, different samples were used for the HPRC (HG00438, HG00733, HG02717, NA20129, HG03453) and HGSVC (HG00731, HG00512, NA19238, NA19650, HG02492). The results are shown in **Figure 3A,** which shows the weighted genotype concordance ^30^ across different types of variants, with the Minigraph-Cactus HPRC graphs showing significantly higher accuracy across all SV variant types than the HGSVC. This improvement can be attributed to the higher quality and number (44 vs 32) of the HPRC vs HGSVC assemblies, as well as the more exact representation of variation, SVs in particular, in the multiple alignment-based Minigraph-Cactus graphs, which would explain the increased delta for SV insertions in particular. This more exact representation also explains why the HPRC graph-based genotypes have fewer very common structural variants (AF>20%) (**Figure 3B),** despite containing significantly more variants (**Figure 3C,D)**. As with the short read variant calling results, the CHM13-based HPRC graph performs generally better than the GRCh38-based graph (**Supplementary Figure 13**)

**Figure 3:**
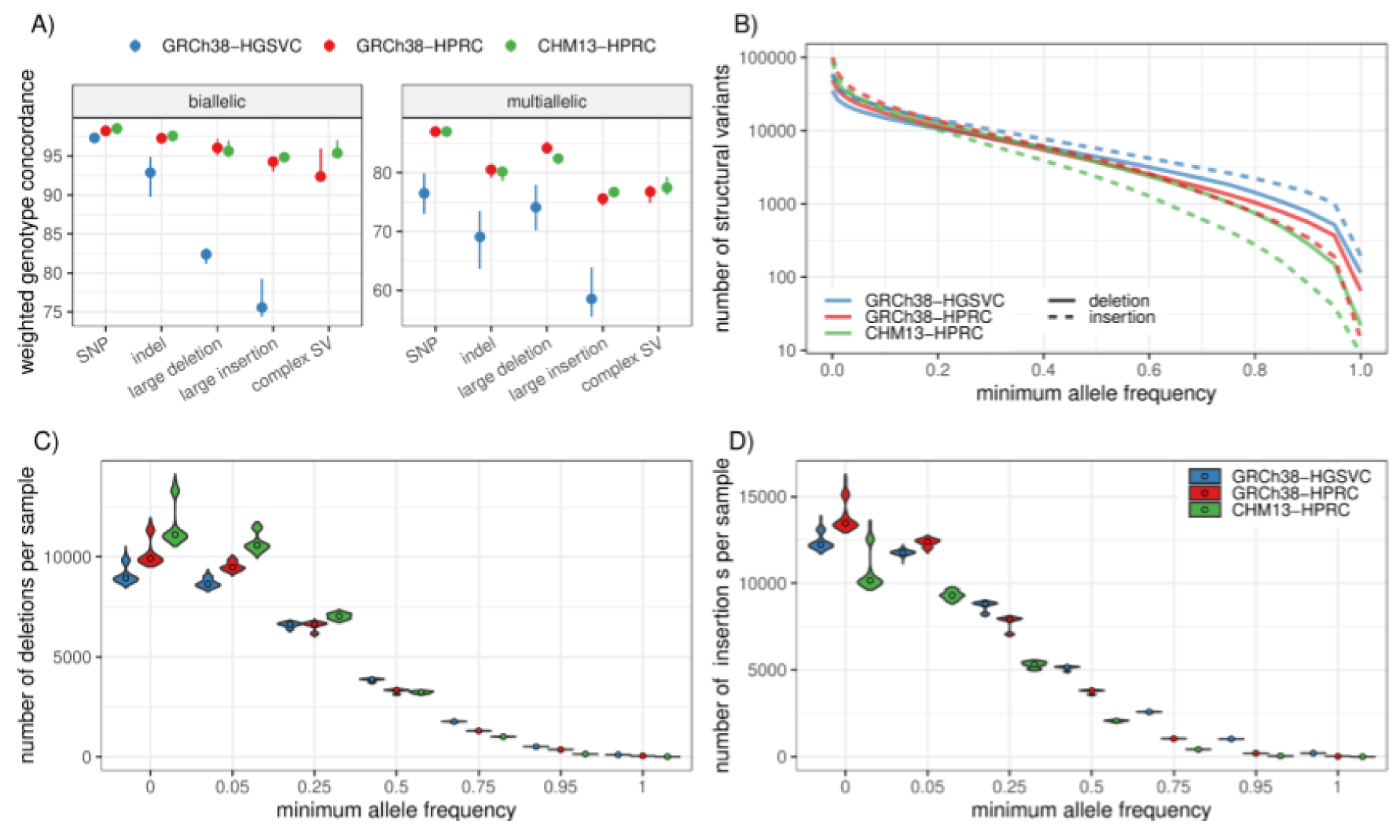
Comparing Pangenome Structural Variant Genotyping. **A)** Leave-one-out PanGenie validation measures the concordance of haplotypes as genotyped by short reads with the haplotypes created via genome assembly. The dots show the medians of five samples independently validated in this way. The lines extend to the minimum and maxiumum values. Note that different samples were used for the HGSVC graph than from the HPRC graphs. **B)** Log-scaled number of structural variants given a minimum allele frequency in the PanGenie genotypes. **C)** The number of SV deletions genotyped per sample, stratified across 6 minimum allele frequency thresholds. The violin plots show the distribution across 368 samples, while the dots represent the median. **D)** The number of SV insertions genotyped per sample, stratified across 6 minimum allele frequency thresholds.

## D. Melanogaster Pangenome

We created a *Drosophila melanogaster* pangenome to demonstrate Minigraph-Cactus’s applicability to non-human organisms. We used 16 assemblies including the reference, dm6 (ISO1), 14 geographically diverse strains described in ^33^, and one additional strain, B7. Their sizes range from 132 to 144 Mb. The allele frequency filtered graph, used for all mapping experiments, was created by removing nodes appearing in < 2 haplotypes leading to a minimum allele frequency of ∼12.5% (compared to 10% in the human graph), and was used only for mapping and genotyping, where private variation in the graph is less helpful. The amount of sequence removed by clipping and filtering is shown in Supplementary Figure 16. The relatively small input meant that we could align it with Progressive Cactus using an all-vs-all (star phylogeny) rather than progressive alignment, and the results are included for comparison. In all, we produced five *D. melanogaster* graphs whose statistics are shown in **Supplementary Table 2**, a process that took roughly 5 hours for the pangenomes (**Supplementary Table 4)** and 19 hours for the progressive Cactus alignments (**Supplementary Table 5**). As in human, adding base-level variants to the SV graph increases its number of nodes and edges by roughly two orders of magnitude. The graph created from the Progressive Cactus alignment has roughly 45% more nodes and edges and over double the total node length (**Supplementary Table 2**).

This is partially explained by the fact that it contains all the sequence filtered out during pangenome construction (**Supplementary Figure 16**) along with interchromosomal alignments. The “core” genome size, which we define as the total length of all nodes present in all samples, of the Minigraph-Cactus pangenome is 110 Mb (**Supplementary Figure 12**, first column), which is roughly half the total size of the graph. This reflects a high diversity among the samples: private transposable element (TE) insertions are known to be abundant in this species ^33^. This diversity is also shown in **Figure 4A**, which graphs the amount of non-reference sequence by the minimum number of samples it is present in, where the private TE insertions would account for much of the nearly 10X differene between the first and second columns. The trend for the number of non-reference nodes is less pronounced (**Supplementary Figure 14**), which implies that the non-reference sequence is accounted for by larger insertion events and smaller variants tend to be more shared. We used the snarl subgraph decomposition ^7^ to compute the variant sites within each graph, i.e. subgraphs equivalent to individual SNPs, indels, SVs, etc. **Supplementary Figure 15** shows the pattern of nesting of the variant sites in the various graphs.

**Figure 4:**
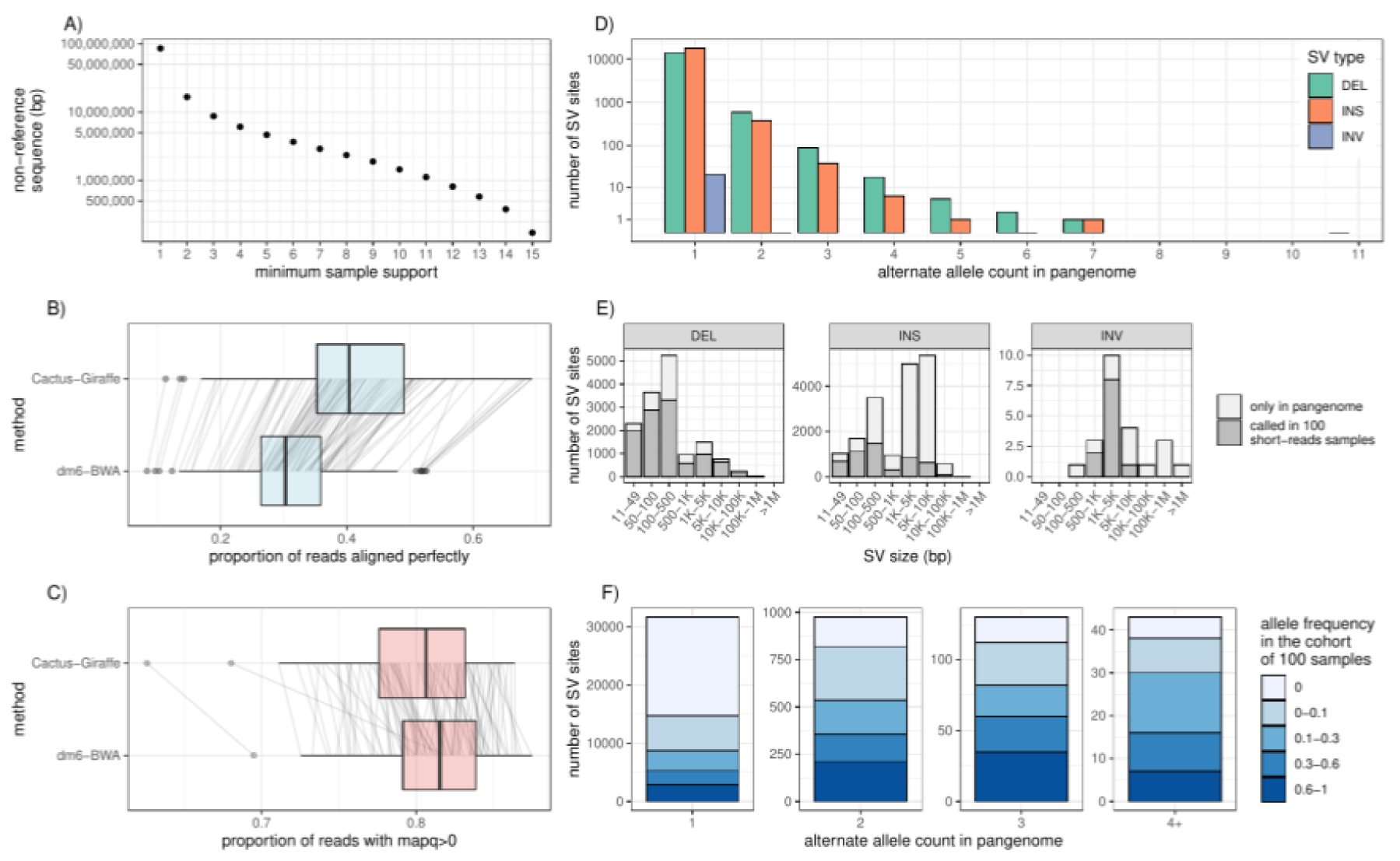
A *Drosophila Melanogaster* Pangenome. **A)** Amount of non-reference sequence by minimum number of haplotypes it occurs in for the *D. melanogaster* pangenome. **B)** Proportion of reads that align perfectly (x-axis) to the filtered pangenome for two approaches (x-axis): “Cactus-Giraffe” where short reads are aligned to the pangenome using vg Giraffe; “dm6-BWA” where reads were mapped to dm6 using BWA-MEM. The boxplots show the median (center line), upper and lower quartiles (box limits), up to 1.5x interquartile range (whiskers), and outliers (points). The lines connect a same sample between the two approaches. **C)** Proportion of reads with a mapping quality above 0. **D)** Distribution of the alternate allele count across each SV site. The x-axis represents the number of assemblies in the pangenome that support a SV. The y-axis is log-scaled. **E)** The size distribution (x-axis) of different SV types (panels). The SV sites are separated in two groups: SV sites that were called in at least one sample from the cohort of 100 samples with short reads (dark grey); SV sites only present in the pangenome (light grey). **F)** Fraction of SVs of different frequency in the cohort of 100 samples (color) compared to their frequency in the pangenome (x-axis).

## Short-read Mapping

The *Drosophila melanogaster* Genetic Reference Panel (DGRP) consists of 205 inbred genomes ^34^, unrelated to the 16 strains used to construct the pangenome. We used short reads from this dataset to evaluate mapping performance for our pangenome graph. We selected 100 samples for our evaluation, filtering the dataset to include only samples with a single SRA accession and Illumina sequencing with >15X coverage. We mapped these samples to the allele frequency filtered pangenome graph with vg giraffe in “fast” mode, and to dm6 using BWA-MEM. We counted the number of mapped reads, reads with perfect alignment, and reads with a mapping quality above 0. We found that the number of reads aligning perfectly drastically increased (**Figure 4B**), with on average 41.1% of the reads aligning perfectly to the pangenome compared to on average 31.0% when aligning reads with BWA on dm6. As in our results in human presented above, we observe a decrease in the number of reads mapped with a mapping quality above 0 when mapping to the pangenome (80.0% vs 81.1% on average, **Figure 4C**).

## Small Variants

We projected pangenomic mappings to dm6, and used FreeBayes ^35^ (in the absence of a high quality DeepVariant model) to call variants on these mappings and those from BWA-MEM (see Methods). We then compared the variant calls that were called by both approaches, and those that were called by only one. While variant sites called by both methods showed similar quality scores, there were more sites unique to our pangenomic approach compared to sites found only by mapping reads to the linear dm6 genome. This increase was observed across different quality thresholds (**Supplementary Figure 17A,C**). Overall, that meant that slightly more variants are called when mapping short reads to the pangenome and projecting them to dm6. For example, on average 740,696 small variants had a quality above 0.1 compared to 738,570 when reads were mapped to the dm6 with BWA-MEM (**Supplementary Figure 17B**). For genotype quality above 10, 705,320 small variants were called versus 700,385 (**Supplementary Figure 17D**). We also noticed a lower rate of heterozygous variants called when mapping the reads to the pangenome first (13.2% vs 18.1% on average per sample, **Supplementary Figure 18**). Due to the high inbreeding of these samples, we expect only a small fraction of variants to truly be segregating ^34^.

## Structural Variants

The variant sites in the pangenome (snarls) were decomposed into canonical structural variants based on the assembly paths in the pangenome (see Methods). In the pangenome, most of the SVs are rare and supported by one or two assemblies (**Figure 4D**). Of note, the known In(3R)C inversion ^36^ is present in the pangenome, along with 23 other smaller inversions. Structural variants were also genotyped from the short read alignments to the pangenome using vg ^5^ (see Methods). Even though the genotyping used short reads and the pangenome was frequency-filtered, 47.8% of the SVs in the pangenome were found when genotyping the 100 samples (on the filtered pangenome) with short-read data. Both the full set of SVs in the pangenome and the subset genotyped from the short read data span the full size spectrum of deletions, insertions and a few inversions (**Figure 4E**). As expected, SVs that were seen in multiple assemblies in the pangenome tended to have higher allele frequencies in the cohort of 100 samples (**Figure 4F**). Both rare and more common SVs spanned the full spectrum of SV size and repeat profile, from the shorter simple repeats and satellite variation to the larger transposable element polymorphisms of LTR/Gypsy, LTR/Pao, and LINE/I-Jockey elements, among others (**Supplementary Figure 19**).

## Discussion

The coordinate system provided by the human reference genome assembly has been vital to nearly all research in human genetics, but it can also be a source of bias. This bias can take the form of unmappable reads in the presence of diverse samples^37^ or, more subtly, variant calls being skewed towards the reference allele^3, 6^. Pangenome graphs have been shown to be effective at reducing reference bias, but their construction has, until now, been limited by trade-offs. Either the graphs needed to be constructed from variant calls against a reference^3, 6, 32^, and therefore unable to properly represent nested variation while still suffering from some reference bias, or they were limited to only structural variants^16^ and unable to effectively be used for short read mapping with current tools^6^. The method we present here overcomes these issues by constructing a pangenome graph directly from a multiple genome alignment that represents nearly all the variation within its inputs.

The challenges of effectively leveraging pangenome graphs for human data do not end at construction. Tooling for analysis, such as read mapping and genotyping, which by definition is more complex for graphs than single reference genomes, is essential. To this end we have ensured that graphs produced with Minigraph-Cactus are the first to be compatible with the majority of state-of-the art pangenome tools (re-engineering the tools as necessary) such as vg^3, 6^, Giraffe^3, 5, 6^, PanGenie^30^ and GraphAligner^27^. These tools are all free and open source. Graphs constructed with Minigraph-Cactus are also freely available for download from the Cactus website and through the HPRC^38^.

To demonstrate the usefulness of these graphs and tools, we showed that Illumina and Hifi reads can be mapped with higher identity and fewer split mappings, respectively, to the pangenome than the linear reference. In the former case, the mappings are used to also improve accuracy of short-read variant calling, and we are hopeful that similar gains will be made with long reads when pangenomics tools for variant calling with them are developed. The representation of structural variants in our multiple-alignment based graphs also show considerable improvements in genotyping accuracy when compared to previous methods that rely on merging reference-based calls.

In the case of DeepVariant and PanGenie, the pangenome graph is used in the context of existing reference-based formats such as BAM and VCF. This allows users to augment their existing workflows with pangenomes with minimal changes, which we think will be key to fostering more widespread adoption of pangenomics methods. Still, such projections back to a linear reference can be lossy, especially in complex regions. While GAF is being increasingly adopted as the standard read mapping format for pangenomes, there is no corresponding graph-based alternative to VCF in use that we are aware of, and the necessity of always projecting variants back to VCF for analysis is a bottleneck to reaching the full potential of pangenome graphs. True graph-based genotyping formats and tools are needed.

Minigraph-Cactus requires at least one chromosome-level input assembly in order to be used as a reference backbone and, in general, the quality and usefulness of the pangenome will increase with the quality and completeness of all the input assemblies. We do not think this will be a bottleneck for most species going forward as it will soon be routine to produce large numbers of reference, or even “telomere-to-telmore” quality genomes for many species due to advances in sequencing technology and assembly tools. In the present work, we have quantified the impact of reference genome assembly quality on our pangenomes and their applications. Even though both GRCh38 and CHM13 are included in all HPRC graphs we constructed, the choice of which to use as a reference backbone influences the topology and completeness of the graph, and in virtually all genome-wide measures of mapping, variant calling and genotyping performance, we found the CHM13-based graph to be superior. In the case of variant calling with Giraffe-DeepVariant, we showed that the CHM13-based graph was able to improve upon the state-of-the art accuracy of the GRCh38-based graph, even when making calls on GRCh38. We therefore think our pangenomes could help some users who would otherwise be reluctant to switch to reference assemblies, still take advantage of them.

Building upon previous work in pangenomics, the HPRC has shown that high-quality genome assemblies can be leveraged to provide a better window into structural variation, as well as to reduce bias incurred by relying on a single reference. The pangenome graph representation has been fundamental to this work, but graph construction remains an active research area. The key challenges stem not just from the computational difficulty of multiple genome alignment, particularly in complex regions, but also from fundamental questions about the tradeoffs between complexity and usability. While developing Minigraph-Cactus, we sought a method to construct graphs with as much variation as possible, while still serving as useful inputs for current pangenome tools like vg and PanGenie.

Some of the compromises made to make our method practical represent exciting challenges for future work in both pangenome construction and applications. Pangenomes from Minigraph-Cactus cannot be used, for instance, to study centromeres. The omission of interchromosomal events will likewise preclude useful cancer pangenomes or studies into acrocentric chromosome evolution ^39^. We are also interested in ways to remove the necessity of filtering the graph to get optimal mapping performance by using an online method at mapping time to identify a subgraph that most closely relates to the reads of a given sample. Progressive Cactus alignments can be combined and updated and, as data sets become larger, this functionality is becoming more necessary for pangenome alignments. Comprehensive tooling to update pangenomes by adding, removing or updating assemblies is an area of future work.

Pangenomics has its origin in non-human species, and as the assembly data becomes available, we will see pangenomes being produced for a wide array of organisms. Already there is data for a number of species, from tomato ^40^ to cow ^41^. In this work, we constructed a *D. melanogaster* pangenome as a proof of concept to show that our method can also be used on other non-human organisms. We hope that others will use the Minigraph-Cactus pipeline to produce useful graphs from sets of genome assemblies for their species of interest. Large-scale alignments are resource intensive, and the 90-human pangenomes required nearly three days to compute on a cluster. As such, we’ve made these alignments publically available through the HPRC and will do the same for future releases.

Reference bias can also affect comparative genomics studies. For example, a genomic region can be of interest to a particular sample, but if that region happens to be missing from the reference genome due intraspecies diversity or assembly errors, it would be absent from any alignments based solely on that reference. Therefore we expect pangenome references to supplant single genome references for intraspecies population genomics studies, we also see this as the future in interspecies comparative genomics studies

## Methods

### Software and Graph Availability

Minigraph-Cactus is included in Cactus, which is released as source, static binaries and Docker images here: https://github.com/ComparativeGenomicsToolkit/cactus/releases. The user guide is here and includes data and instructions to build a yeast and HPRC pangenome: https://github.com/ComparativeGenomicsToolkit/cactus/blob/master/doc/pangenome.md. Links to the human and *D. melanogaster* pangenome graphs and indexes, as well as those for some other species can be found here: https://github.com/ComparativeGenomicsToolkit/cactus/tree/master/doc/mc-pangenomes/README.md. Please consult https://github.com/ComparativeGenomicsToolkit/cactus/blob/master/doc/mc-paper/README.md for command lines and scripts used for this work. Pangenome Graphs created with our method that were released as part of the HPRC can also be found on the latter’s data portal: https://github.com/human-pangenomics/hpp_pangenome_resources/

### HPRC Graph Construction

The HPRC v1.0 graphs discussed here were created by an older version of the pipeline described above, with the main difference being that the satellite sequence was first removed from the input with dna-brnn ^42^. This procedure is described in detail in ^25^. The amount of sequence removed from the graph, and the reason it was removed, is shown in **Supplementary Figure 2**. Roughly 200 Mb per assembly was excluded, the majority of which was flagged as centromeric (HSat2 or alpha satellite) by dna-brnn ^42^. The “unassigned”, “minigraph-gap” and “clipped” categories denote the sequence that, respectively, did not map well enough to any one chromosome to be assigned to it, intervals > 100kb that did not map with minigraph, and intervals > 10kb that did not align with Cactus. Simply removing all sequence ≥10kb that does not align with Cactus, as described in the methods above, amounts to nearly the same amount of sequence excluded (**Supplementary Figure 3**). The 10kb threshold was used for clipping because it was sufficient to remove all centromeres (as previously identified) with dna-brnn and also because it corresponds to the maximum length of an alignment that can be computed with abPOA. The exact commands to build HPRC graphs referred to in this figure are available here: https://github.com/ComparativeGenomicsToolkit/cactus/blob/91bdd83728c8cdef8c34243f0a52b28d85711bcf/doc/pangenome.md#hprc-graph. They were run using the same Cactus commit: 91bdd83728c8cdef8c34243f0a52b28d85711bcf.

### Filtering Minigraph Mappings and Chromosome Decomposition

Input contigs were labeled “unassigned” above if they could not be confidently mapped to a single reference chromosome during the Minigraph contig mapping phase of the pipeline. For a given contig, this determination was made by identifying the chromosome in the SV graph to which the highest fraction of its bases mapped with exact matches. If this highest fraction was at least three times higher than the second highest, and greater than or equal to a minimum threshold, the contig was assigned to that chromosome, otherwise it was left unassigned (and omitted from the graph). The minimum threshold for chromosome assignment was 75% for contigs with length ≤ 100 kb, 50% for contigs with length in the range (100 kb, 1 Mb] and 25% for with length > 1 Mb. These values were chosen after empirical experimentation specifically to filter out spurious mappings as determined by VCF-based comparison with HiFi-based DeepVariant calls^25^. Contigs filtered in this way are predominantly centromeric (and can’t be confidently mapped anywhere) or small fragments of acrocentric chromosome short arms or segmental duplications without enough flanking sequence to be correctly placed, or regions enriched for putative misjoins (which also occur predominantly within the acrocentric chromosome short arms) ^25^. Such filtering is not needed on chromosome level assemblies.

Despite this filtering process, we found a small number of small contigs that, due to either misassembly or misalignment, confidently map across entire chromosome arms (one of the contig maps near the centromere and the other near the telomere). The chromosome arm-spanning edges introduced by such mappings introduce topological complexities that can hinder downstream tools (for example, all variants on the spanned arm would be considered nested within a large deletion). To prevent this, any mapping that would introduce a deletion edge of 10 Mb or more (tunable by a parameter) relative to the reference path is removed.

Finally, in rare cases, minigraph can map the same portion of a query contig to different target regions in the graph. When manually inspecting these cases, we found that they could lead to spurious variants in the graph when, as above, compared to variant calls directly from HiFi-based DeepVariant calls (Liao et al., 2022). To mitigate these cases, we remove any aligned query interval (pairwise alignments are represented in terms of the query intervals, positions on the contig, and target intervals, paths within the graph) that overlaps another by at least 25% of its length, and whose mapping quality and/or block length is 5X lower than those of the other interval.

### POA-based Cactus Base Aligner

We replaced the base-level alignment refinement (BAR) algorithm that is used to create alignments between the interstitial sequences after the initial anchoring process^20^. Briefly, the original algorithm has two stages. Firstly, from the end of each alignment anchor (termed a block, and defined by a gapless alignment of substrings of the input) it creates a MSA of the unaligned sequences incident with the anchor. Each such MSA has the property that the sequence alignment is pinned from the anchor point, but because of rearrangement, the MSA is not necessarily global, i.e. at the other end of the MSA from the starting anchor point the different sequences may be non-homologous due to genome rearrangement. Secondly, the set of MSAs produced by the first step are refined by a greedy process which seeks to make the set of MSAs, which may overlap in terms of sequence positions, consistent, so resolving, at base-level resolution, the breakpoints of genome rearrangements. For details of this process see the original paper^20^.

The replacement BAR algorithm achieved two things. Firstly, we changed the process in the first step to create MSAs to use the abPOA MSA algorithm^43^. The previous algorithm was based upon the original Pecan MSA process, and scaled quadratically with sequence number, in contrast the new MSA process scales linearly and is overall faster even for small numbers of sequences. In this process we updated abPOA to use the LASTZ default scoring parameters^12^, with the addition of a “long” gap state not used by LASTZ but included within abPOA. Gap parameters were thus: short-gap-open: 400, short-gap-extend: 30, long-gap-open:1200, long-gap-extend:1. Parameters for the long-gap-state were determined by empirical experimentation. Secondly, we fully reimplemented the second step of the BAR algorithm, making it both faster and removing various unnecessary bottlenecks which previously scaled superlinearly but which now all scale linearly with sequence number and length. Importantly, this process did not materially affect the resulting alignments, as judged by extensive unit-and system-level testing.

### Conversion from Multiple Alignment to Sequence Graph

Cactus natively uses Hierarchical Alignment (HAL) format^21^. We developed hal2vg, which converts HAL files to vg formats. It works for both Progressive and Minigraph-Cactus. It works in memory and, for large alignments, is reliant on having chromosomal decomposition of the HAL and simple topology to run efficiently. hal2vg begins by visiting the pairwise alignments in breadth-first order from the root of the underlying guide tree. Contiguous runs of exact matches in the pairwise are “pinched” together to form nodes of a sequence graph using Cactus^15^, and the assemblies themselves are added as “threads” to this graph. SNPs are stored in an auxiliary data structure and used to pinch together transitive exact matches as they arise. For example, if the pairwise alignments of a column (in the multiple alignment) are A->C and C->A, this structure will ensure that the two A’s are pinched together in the sequence graph (which, by definition, only represents exact matches within its nodes). Seqwish^44^ is a recent tool that also induces sequence graphs from sets of pairwise alignments but, because it does not transitively process SNPs in this way, will not work on tree-based sets of pairwise alignments as represented by HAL. Finally, once the sequence graph has been created in memory, it is serialized to disk, path by path, using libbdsg^45^, an API for reading and writing sequence graphs in an efficient, VG-compatible binary format.

### Conversion from Sequence Graph to VCF

By default, all graphs are output in GFA (v1.1), as well as the vg-native indexes: xg, snarls and GBWT formats ^45, 46^. Since VCF remains more widely-supported than these formats, we implemented a VCF exporter in vg (`vg deconstruct)` that is run as part of the Minigraph-Cacatus pipeline. It outputs a site for each snarl in the graph. It uses the haplotype index (GBWT) to enumerate all haplotypes that traverse the site, which allows it to compute phased genotypes. For each allele, the corresponding path through the graph is stored in the AT (Allele Traversal) tag. Snarls can be nested, and this information is specified in the LV (Level) and PS (Parent Snarl) tags, which needs to be taken into account when interpreting the VCF. Any phasing information in the input assemblies is preserved in the VCF.

### HPRC Graph Mapping and Variant Calling

We used 30x Illumina NovaSeq PCR-free short read data HG001, HG002, and HG005, available at gs://deepvariant/benchmarking/fastq/wgs_pcr_free/30x/. The reads were mapped to the pangenome using vg giraffe (v1.37.0). The same reads were mapped to GRCh38 with decoy sequences, but no ALTs using BWA-MEM (v0.7.17). To provide additional baselines, reads were also mapped with vg giraffe to linear pangenomes, i.e. pangenomes containing only the reference genome (GRCh38 or CHM13). The number of reads mapped with different mapping quality (or aligning perfectly) were extracted from the graph alignment file (GAF/GAM files) produced by vg giraffe and from the BAM files produced by BWA-MEM.

Variants were called using the approach described in ^25^. Briefly, the graph alignments were projected to the chromosomal paths (chr 1-22, X, Y) of GRCh38 using vg surject. Once sorted with samtools (v1.3.1), the reads were realigned using bamleftalign (Freebayes v1.2.0) ^35^ and ABRA (v2.23) ^47^. DeepVariant (v1.3) ^28^ then called small variants using models trained for the HPRC pangenome ^25^. We used the same approach when calling small variants using the CHM13-based pangenome and when projecting to CHM13 chromosomal paths.

### Evaluation of small variant calls

Calls on GRCh38 were evaluated as in ^25^, i.e. using the Genome In A Bottle (GIAB) benchmark and confident regions for each of the three samples ^48^. For HG002, the Challenging Medically Relevant Genes (CMRG) truth set v1.0 ^29^ was also used to evaluate small variants calls in those challenging regions. The evaluation was performed by hap.py ^49^ v0.3.12 via the jmcdani20/hap.py:v0.3.12 docker image. When evaluating calls made against the GRCh38 chromosomal paths using the CHM13-based pangenome, we excluded regions annotated as false-duplications and collapsed in GRCh38.

These regions do not have a well-defined truth label in the context of CHM13. We used the “GRCh38_collapsed_duplication_FP_regions”, “GRCh38_false_duplications_correct_copy”, “GRCh38_false_duplications_incorrect_copy”, and “GRCh38_population_CNV_FP_regions” region sets available at https://github.com/genome-in-a-bottle/genome-stratifications.

To evaluate the calls made on CHM13 v1.1, we used two approaches. First, the calls from CHM13 v1.1 were lifted to GRCh38 and evaluated using the GRCh38 truth sets described above (GIAB v4.2.1 and CMRG v1.0). For this evaluation, we also lifted these GRCh38-based truth sets to CHM13 v1.1 to identify which variants of the truth set are not visible on CHM13 because they are homozygous for the CHM13 reference allele. Indeed, being homozygous for the reference allele, those calls will not be present in the VCF because there are no alternate alleles to find. These variants were excluded from the truth set during evaluation. The second approach was to evaluate the calls in CHM13 v1.1 directly. To be able to use the CMRG v1.0 truth set provided by the GIAB, we lifted the variants and confident regions from CHM13 v1.0 to CHM13 v1.1. The CMRG v1.0 truth set focuses on challenging regions, but still provides variant calls across the whole genome. Hence, we used those variants to evaluate the performance genome-wide although restricting to a set of confident regions constructed by intersecting the confident regions for HG002 from GIAB v4.2.1 (lifted from GRCh38 to CHM13 v1.1), and the alignment regions produced by dipcall in the making of the CMRG v1.0 truth set (https://ftp-trace.ncbi.nlm.nih.gov/ReferenceSamples/giab/release/AshkenazimTrio/HG002_NA2 4385_son/CMRG_v1.00/CHM13v1.0/SupplementaryFiles/HG002v11-align2-CHM13v1.0/HG002 v11-align2-CHM13v1.0.dip.bed). Finally, we used the preliminary HG002 truth set from GIAB on CHM13 v2.0 which is equivalent to CHM13 v1.1 with the added chromosome Y from HG002. The calls in this set wer based on aligning a high-confidence assembly using dipcall ^50^ (labeled in figure as “dipcall CHM13 v2.0”). Here again, we intersected the confident regions with the GIAB v4.2.1 confident regions lifted from GRCh38 to CHM13.

In all experiments described above, the variants (VCF files) were lifted over using Picard (v2.27.4) ^51^ LiftoverVcf and the RECOVER_SWAPPED_REF_ALT option. Regions (BED files) were lifted with liftOver ^52^.

Finally, we compared in greater detail the calling performance using the GRCh38-based and CHM13-based pangenomes by stratifying the evaluation across genomic region sets provided by the GIAB (https://github.com/genome-in-a-bottle/genome-stratifications). These regions included, for example, different types of challenging regions like segmental duplications, simple repeats, transposable elements.

### Alignment of long reads

HiFi reads from HG002, HG003, and HG004 were downloaded from Genome in a Bottle FTP site, ftp-trace.ncbi.nlm.nih.gov.: /giab/ftp/data/AshkenazimTrio/HG002_NA24385_son/PacBio_CCS_15kb_20kb_chemistry2/reads/m64011_190830_220126.fastq.gz /giab/ftp/data/AshkenazimTrio/HG003_NA24149_father/PacBio_CCS_15kb_20kb_chemistry2/reads/PBmixSequel729_1_A01_PBTH_30hours_19kbV2PD_70pM_HumanHG003.fastq.gz /giab/ftp/data/AshkenazimTrio/HG004_NA24143_mother/PacBio_CCS_15kb_20kb_chemistry2/ uBAMs/m64017_191115_211223.hifi_reads.bam The reads were then aligned to the pangenomes (after being converted to fastq with samtools fastq in the case of HG004) using GraphAligner (v.0.13) with ‘-x vg’ with .gam output.. We parsed the GAM output to extract the first record as primary alignment. By overlapping the other alignment records with the primary alignment, we identified reads with split-mapping, i.e. with part of the read is mapped to a different location from the primary alignment. The alignment identity is reported by GraphAligner and was also extracted from the GAM.

### SV Genotyping with PanGenie

Variants corresponding to nested sites in the HPGRC graph-derived VCFs were decomposed as described in^25^ before running PanGenie version v2.1.0 with its default parameters. The HGSVC v.4.0 “lenient set”^32^ was also included, but did not require decomposition. These three VCFs, annotated with all computed genotypes, are available for download here: https://zenodo.org/record/7669083. The genotyped samples were chosen by randomly selecting 100 trios for the 1KG data, 20 from each superpopulation. Samples present in HPRC and HGSVC were also included, for a total of 368. High coverage short reads from the 1KG^31^ were used for genotyping. The leave-one-out experiments were performed as described in ^25^ and, like in that work, variants were “collapsed” using `truvari collapse -r 500 -p 0.95 -P 0.95 -s 50 -S 100000` from Truvari^53^ version 3.5.0 when comparing counts of genotyped variants (**Figure 3 B,C,D**). This is because near-identical insertions in the graph become completely separate variants in the VCF when, for the purposes of this comparison, we wish to treat them the same. SV deletions (insertions) were sites with reference alleles of length >= 50 (1) and alt alleles of length 1 (>=50). Sites that did not meet this criteria but had a reference or alt allele of length >= 50 were classified as “SV Other”.

### *D. Melanogaster* Graph Construction

The *D. Melanogaster* pangenome was created using Minigraph-Cactus using the procedure described in The Minigraph-Cactus Pangenome Pipeline secion . Progressive Cactus was run on the same input (which implies a star phylogeny) and was exported to vg with `hal2vg`.

### *D. Melanogaster* Variant Decomposition

The variant sites in the pangenome (snarls, aka bubbles) were decomposed into canonical structural variants using a script developed for the HPRC analysis ^25^. In brief, each allele in the deconstructed VCF specifies the corresponding path in the pangenome. The script follows these paths and, comparing them with the dm6 reference path, enumerates each canonical variant (SNP, indels, structural variants). The frequency of each variant in the pangenome corresponds to the number of assemblies that traverse their paths.

### *D. Melanogaster* Graph Mapping and Variant Calling

The DGPR samples used are listed in **Supplementary Table 6**. Short reads were obtained using `fasterq-dump –split 3` on the accessions in the last column of this table. Each read pair was mapped to the allele-frequency filtered graph with `vg giraffe` and to dm6 with BWA-MEM.

`vg call` was used to to genotype variants in the pangenome. For each sample, these variant calls were decomposed into canonical SVs using the same approach described above on the HPRC deconstructed VCF. The SV calls were then compared to the SVs in the pangenome using the sveval package ^5^ which matches SVs based on their types, sizes and location.Because SVs are genotyped using the same pangenome, they are expected to be relatively similar, and we can use standard “collapse” criteria to cluster them in SV sites. Two SVs were matched if: their regions had a reciprocal overlap of at least 90% for deletions and inversions; they were located at less than 100bp from each other, and their inserted sequences were at least 90% similar for insertions. The same approach was used to cluster the SVs alleles into the SV sites reported in the text and figures. The SV alleles were annotated with RepeatMasker (v4.0.9). We assigned a repeat class to a SV if more than 80% of the allelic sequence was annotated as such. The 80% threshold was chosen by inspecting the distribution and observing a negligible number of events below this value.

We used `vg surject` to produce BAM files referenced on dm6 from the mappings to the pangenome, and FreeBayes v1.3.6 ^35^ (in the absence of a high quality DeepVariant model) to call variants on these mappings and those from BWA-MEM. Single-sample VCFs were merged with `bcftools merge.`

To compare the variant calls by both approaches, we used bcftools ^54^ (v1.10.2) to normalize the VCFs (bcftools norm), and compare them (bcftools isec) to mark variant sites where both approaches call a variant, and sites where only one approach does. We compared the number of calls in each category, across samples, and for different minimum variant quality thresholds (QUAL field or genotype quality GQ field).

## Acknowledgements

We thank Anthony D. Long for many suggestions and insights regarding the *D. Melanogaster* data, and the whole vg team for their work to create and maintain vg upon which much of this work depends.

## Human Pangenome Reference Consortium Authorship

Haley J. Abel^1^,Lucinda L Antonacci-Fulton^2^, Mobin Asri^3^, Gunjan Baid^4^, Carl A. Baker^5^, Anastasiya Belyaeva^4^, Konstantinos Billis^6^, Guillaume Bourque^7,8,9^, Silvia Buonaiuto^1^, Andrew Carroll^4^, Mark JP Chaisson^11^, Pi-Chuan Chang^4^, Xian H. Chang^3^, Haoyu Cheng^12,13^, Justin Chu^12^, Sarah Cody^2^, Vincenza Colonna^10,14^, Daniel E. Cook^4^, Robert M. Cook-Deegan^15^, Omar E. Cornejo^16^, Mark Diekhans^3^, Daniel Doerr^17^, Peter Ebert^17^, Evan E. Eichler^5,18^, Jordan M. Eizenga^3^, Susan Fairley^6^, Olivier Fedrigo^19^, Adam L. Felsenfeld^20^, Xiaowen Feng^12,13^, Christian Fischer^14^, Paul Flicek^6^, Giulio Formenti^19^, Adam Frankish^6^, Robert S. Fulton^2^, Shilpa Garg^22^, Erik Garrison^14^, Carlos Garcia Giron^6^, Richard E. Green^23,24^, Cristian Groza^25^, Andrea Guarracino^26^, Leanne Haggerty^6^, Ira Hall^27,28^, William T Harvey^5^, Marina Haukness^3^, David Haussler^3,18^, Simon Heumos^29,30^, Glenn Hickey^3^, Kendra Hoekzema^5^, Thibaut Hourlier^6^, Kerstin Howe^31^, Miten Jain^32^, Erich D. Jarvis^33,18^, Hanlee P. Ji^34^, Alexey Kolesnikov^4^, Jan O. Korbel^35^, Jennifer Kordosky^5^, Sergey Koren^36^, HoJoon Lee^34^, Alexandra P. Lewis^5^, Heng Li^12,13^, Wen-Wei Liao^2,37,27^, Shuangjia Lu^27^, Tsung-Yu Lu^38^, Julian K. Lucas^3^, Hugo Magalhães^17^, Santiago Marco-Sola^39,40^, Pierre Marijon^17^, Charles Markello^3^, Fergal J. Martin^6^, Ann McCartney^36^, Jennifer McDaniel^41^, Karen H. Miga^3^, Matthew W. Mitchell^42^, Jean Monlong^3^, Jacquelyn Mountcastle^19^, Katherine M. Munson^5^, Moses Njagi Mwaniki^43^, Maria Nattestad^4^, Adam M. Novak^3^, Sergey Nurk^36^, Hugh E. Olsen^3^, Nathan D. Olson^41^, Benedict Paten^3^, Trevor Pesout^3^, Adam M. Phillippy^36^, Alice B. Popejoy^44^, David Porubsky^5^, Pjotr Prins^14^, Daniela Puiu^45^, Mikko Rautiainen^36^, Allison A Regier^2^, Arang Rhie^36^, Samuel Sacco^46^, Ashley D. Sanders^47^, Valerie A. Schneider^48^, Baergen I. Schultz^20^, Kishwar Shafin^4^, Jonas A. Sibbesen^49^, Jouni Sirén^3^, Michael W. Smith^20^, Heidi J. Sofia^20^, Ahmad N. Abou Tayoun^50,51^, Françoise Thibaud-Nissen^48^, Chad Tomlinson^2^, Francesca Floriana Tricomi^6^, Flavia Villani^14^, Mitchell R. Vollger^5,52^, Justin Wagner^41^, Brian Walenz^36^, Ting Wang^53^, Jonathan M. D. Wood^31^, Aleksey V. Zimin^45,54^, Justin M. Zook^41^

1 Division of Oncology, Department of Internal Medicine, Washington University School of Medicine, St. Louis, MO 63110, USA

2 McDonnell Genome Institute, Washington University School of Medicine, St. Louis, MO 63108, USA

3 UC Santa Cruz Genomics Institute, University of California, Santa Cruz, 1156 High St, Santa Cruz, CA, USA

4 Google LLC, 1600 Amphitheater Pkwy, Mountain View, CA 94043, USA

5 Department of Genome Sciences, University of Washington School of Medicine, Seattle, WA 98195, USA

6 European Molecular Biology Laboratory, European Bioinformatics Institute, Wellcome Genome Campus, Cambridge, CB10 1SD, UK

7 Department of Human Genetics, McGill University, Montreal, Québec H3A 0C7, Canada

8 Canadian Center for Computational Genomics, McGill University, Montreal, Québec H3A 0G1, Canada 9 Institute for the Advanced Study of Human Biology (WPI-ASHBi), Kyoto University, Kyoto 606-8501, Japan

10 Institute of Genetics and Biophysics, National Research Council, Naples 80111, Italy

11 University of Southern California, Quantitative and Computational Biology, Los Angeles, CA, USA 12 Department of Data Sciences, Dana-Farber Cancer Institute, Boston, MA 02215, USA

13 Department of Biomedical Informatics, Harvard Medical School, Boston, MA 02215, USA

14 Department of Genetics, Genomics and Informatics, University of Tennessee Health Science Center, Memphis, TN 38163, USA

15 Arizona State University, Barrett & O’Connor Washington Center, Washington DC, USA 16 School of Biological Sciences, Washington State University, Pullman WA 99163, USA

17 Institute for Medical Biometry and Bioinformatics, Medical Faculty, Heinrich Heine University Düsseldorf, Düsseldorf, Germany

18 Howard Hughes Medical Institute, Chevy Chase, MD 20815, USA

19 The Vertebrate Genome Laboratory, The Rockefeller University, New York, NY 10065, USA

20 National Institutes of Health (NIH)–National Human Genome Research Institute, Bethesda, MD, USA 21 Center for Computational and Genomic Medicine, The Children’s Hospital of Philadelphia, Philadelphia, PA 19104, USA.

22 Department of Biology, University of Copenhagen, Denmark

23 Department of Biomolecular Engineering, University of California, Santa Cruz, 1156 High St., Santa Cruz, CA 95064, USA

24 Dovetail Genomics, Scotts Valley, CA 95066, USA

25 Quantitative Life Sciences, McGill University, Montreal, Québec H3A 0C7, Canada 26 Genomics Research Centre, Human Technopole, Milan 20157, Italy

27 Department of Genetics, Yale University School of Medicine, New Haven, CT 06510, USA 28 Center for Genomic Health, Yale University School of Medicine, New Haven, CT 06510, USA 29 Quantitative Biology Center (QBiC), University of Tübingen, Tübingen 72076, Germany

30 Biomedical Data Science, Department of Computer Science, University of Tübingen, Tübingen 72076, Germany

31 Tree of Life, Wellcome Sanger Institute, Hinxton, Cambridge, CB10 1SA, UK 32 Northeastern University, Boston, MA 02115, USA

33 The Rockefeller University, New York, NY 10065, USA

34 Division of Oncology, Department of Medicine, Stanford University School of Medicine, Stanford, CA, 94305, USA

35 European Molecular Biology Laboratory, Genome Biology Unit, Meyerhofstr. 1, 69117 Heidelberg, Germany

36 Genome Informatics Section, Computational and Statistical Genomics Branch, National Human Genome Research Institute, National Institutes of Health, Bethesda, MD 20892, USA

37 Department of Medicine, Washington University School of Medicine, St. Louis, MO 63110, USA

38 University of Southern California, Quantitative and Computational Biology, 3551 Trousdale, Pkwy, Los Angeles, CA, USA

39 Computer Sciences Department, Barcelona Supercomputing Center, Barcelona, Spain

40 Departament d’Arquitectura de Computadors i Sistemes Operatius, Universitat Autònoma de Barcelona, Barcelona, Spain

41 Material Measurement Laboratory, National Institute of Standards and Technology, Gaithersburg, MD 20877, USA

42 Coriell Institute for Medical Research, Camden, NJ 08103, USA

43 Department of Computer Science, University of Pisa, Pisa 56127, Italy

44 Department of Public Health Sciences, University of California, Davis, One Shields Avenue, Medical Sciences 1C, Davis, CA 95616

45 Department of Biomedical Engineering, Johns Hopkins University, Baltimore 21218, MD, USA

46 Department of Ecology & Evolutionary Biology, University of California, Santa Cruz, 1156 High St, Santa Cruz, CA, USA

47 Berlin Institute for Medical Systems Biology, Max Delbrück Center for Molecular Medicine in the Helmholtz Association, Berlin, Germany

48 National Center for Biotechnology Information, National Library of Medicine, National Institutes of Health, Bethesda, MD 20894, USA

49 Center for Health Data Science, University of Copenhagen, Denmark

50 Al Jalila Genomics Center of Excellence, Al Jalila Children’s Specialty Hospital, Dubai, UAE

51 Center for Genomic Discovery, Mohammed Bin Rashid University of Medicine and Health Sciences, Dubai, UAE

52 Division of Medical Genetics, University of Washington School of Medicine, Seattle, WA 98195, USA 53 Department of Genetics, Washington University School of Medicine, St. Louis, MO 63110, USA

54 Center for Computational Biology, Johns Hopkins University, Baltimore, MD 21218, USA

## Supplement

**Supplementary Figure 1:**
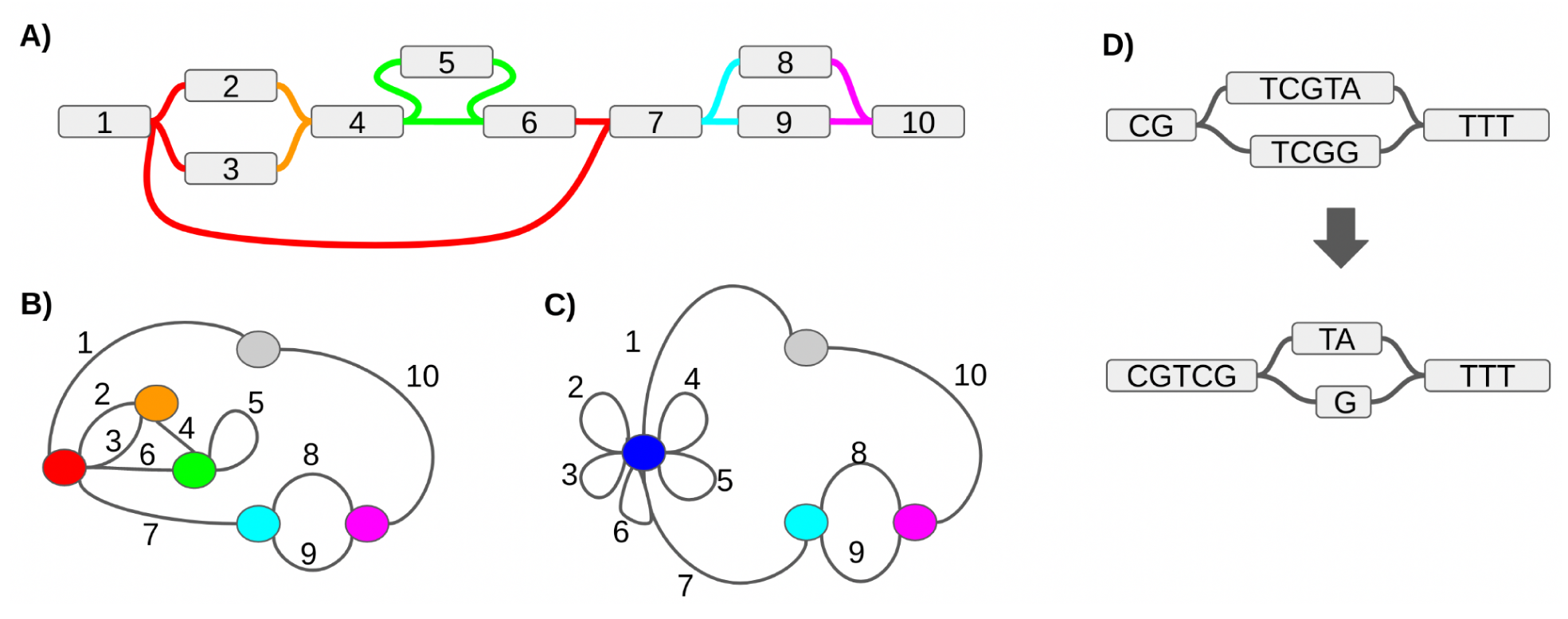
A-C: Cactus graph eample. **A)** A sequence graph where the ID of each node is shown but not its sequence. We consider two edges “connected” if they are incident to the same “end” of the same node. Connected components of edges under this definition are grouped together, with each component being given a separate colour. **B)** Each connected component of edges in the sequence graph is grouped together into a node. Each node in the sequence graph is transformed into an edge. A “root” node (gray) is created to connect to all node ends that have degree 0 in the sequence graph (stubs). **C)** The cactus graph is created by merging together all 3-edge-connected components of nodes in the graph from B). This graph has the property that no edge is part of more than one simple cycle. **D)** 3bp of redundant sequence, “TCG”, is removed with GFAffix. This sequence is redundant in the sense that its removal does not affect the number of possible haplotype paths through the graph.

**Supplementary Figure 2:**
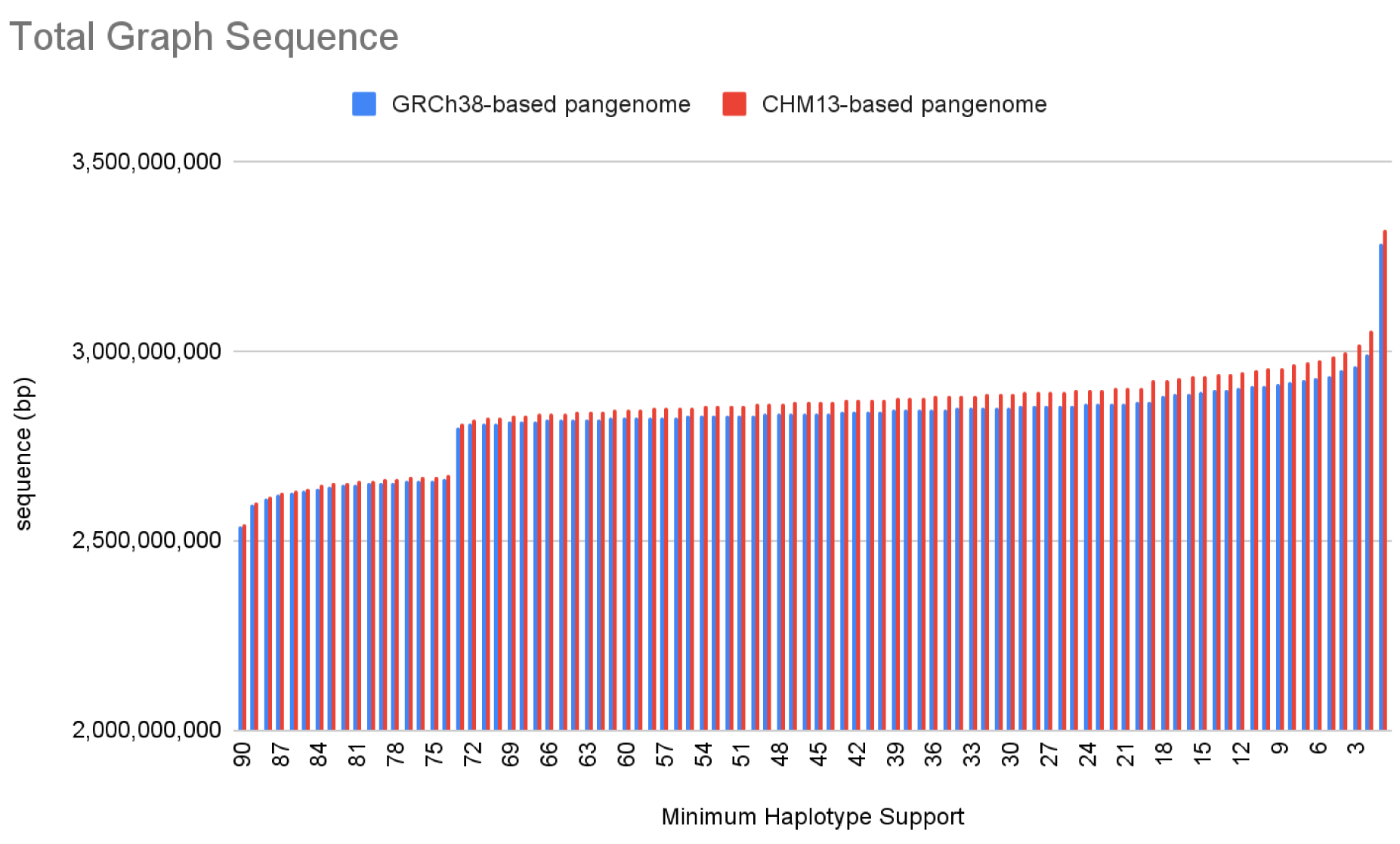
The amount of sequence in the HPRC graphs by the minimum number of haplotypes that contain it. The step in the graph is due to 14 male haplotypes not possessing an X chromosome.

**Supplementary Figure 3:**
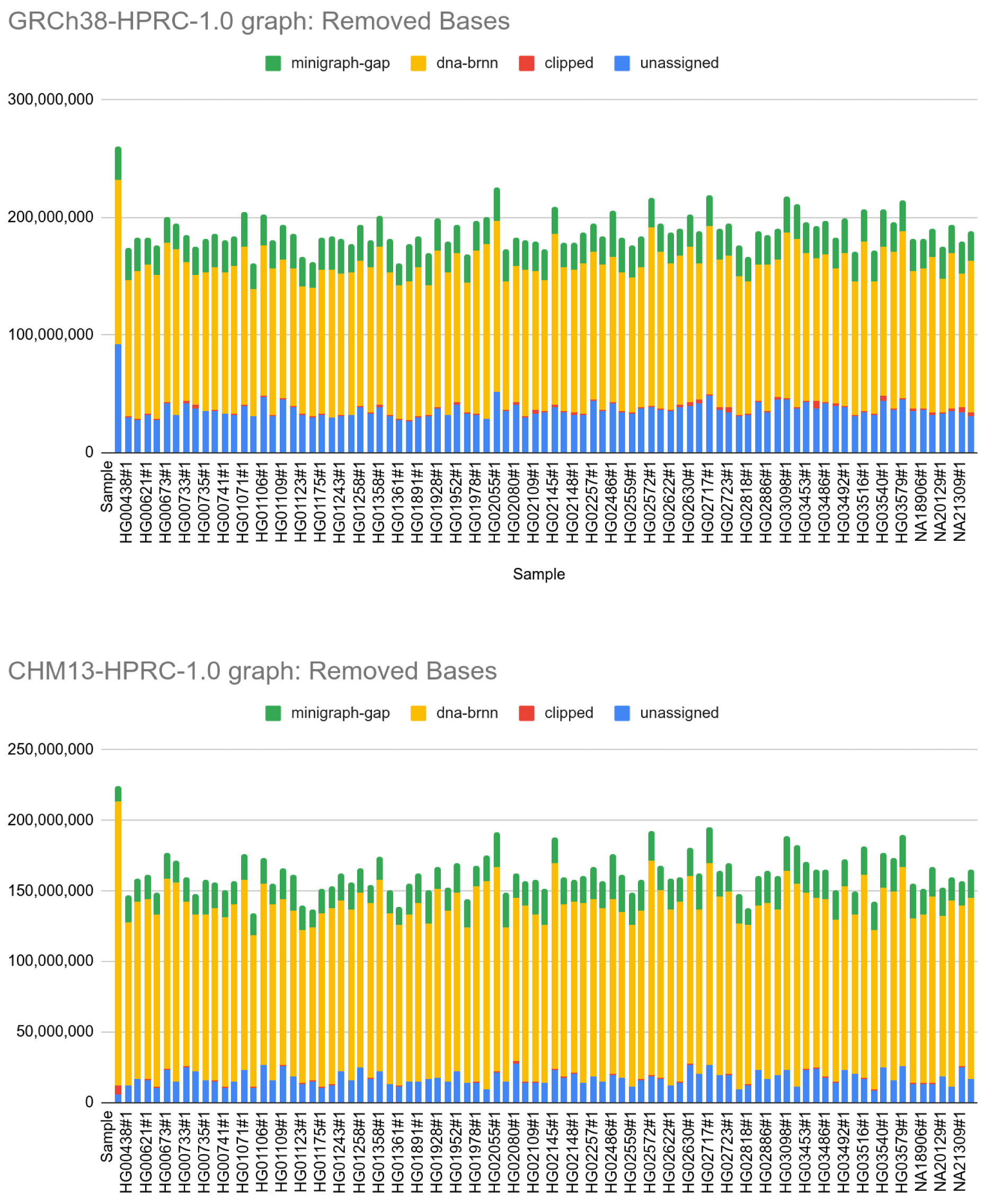
Sequence excluded from the HPRC pangenomes.

**Supplementary Figure 4:**
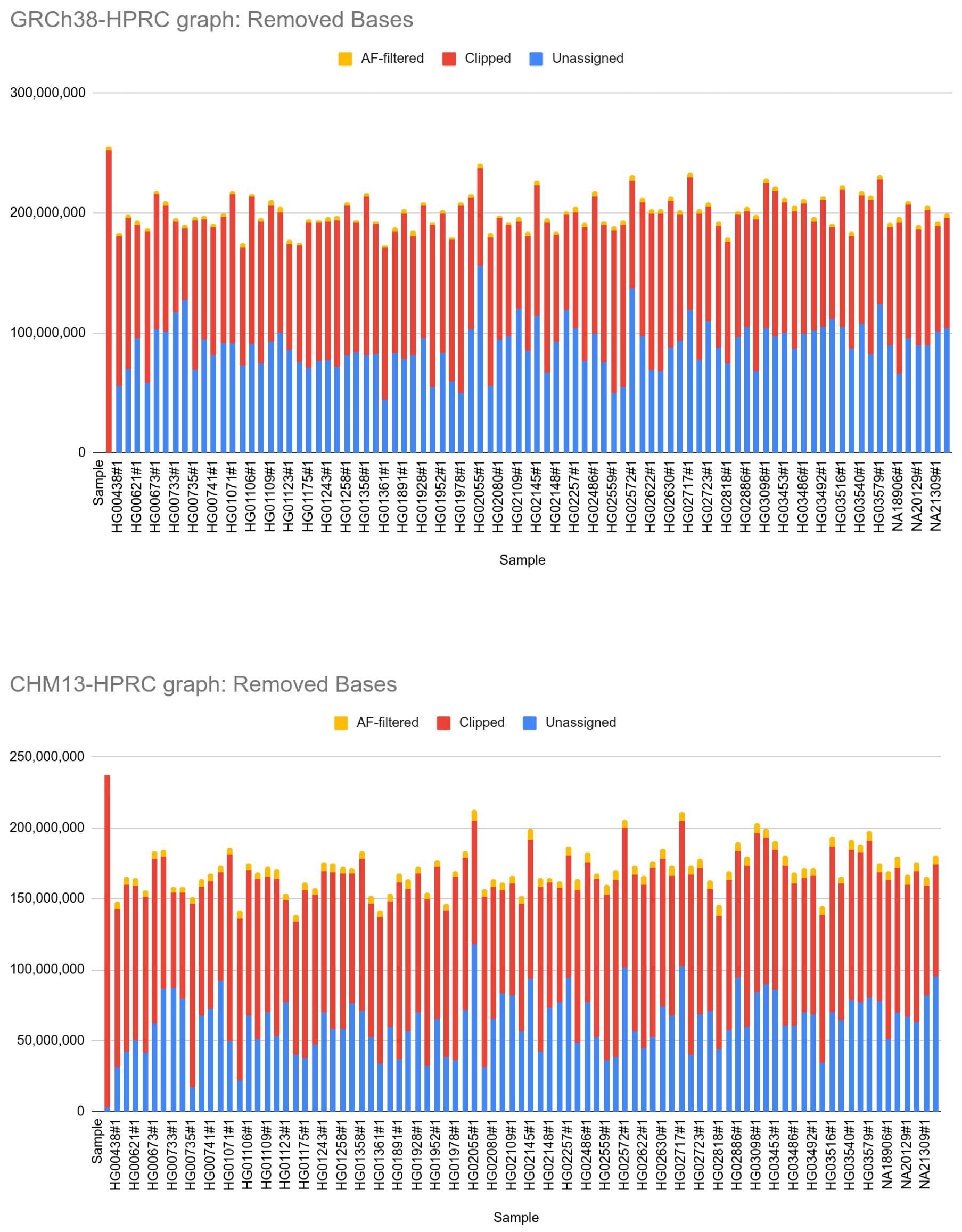
Sequence excluded from the HPRC pangenomes when using the current pipeline (without dna-brnn preprocessing).

**Supplementary Figure 5:**
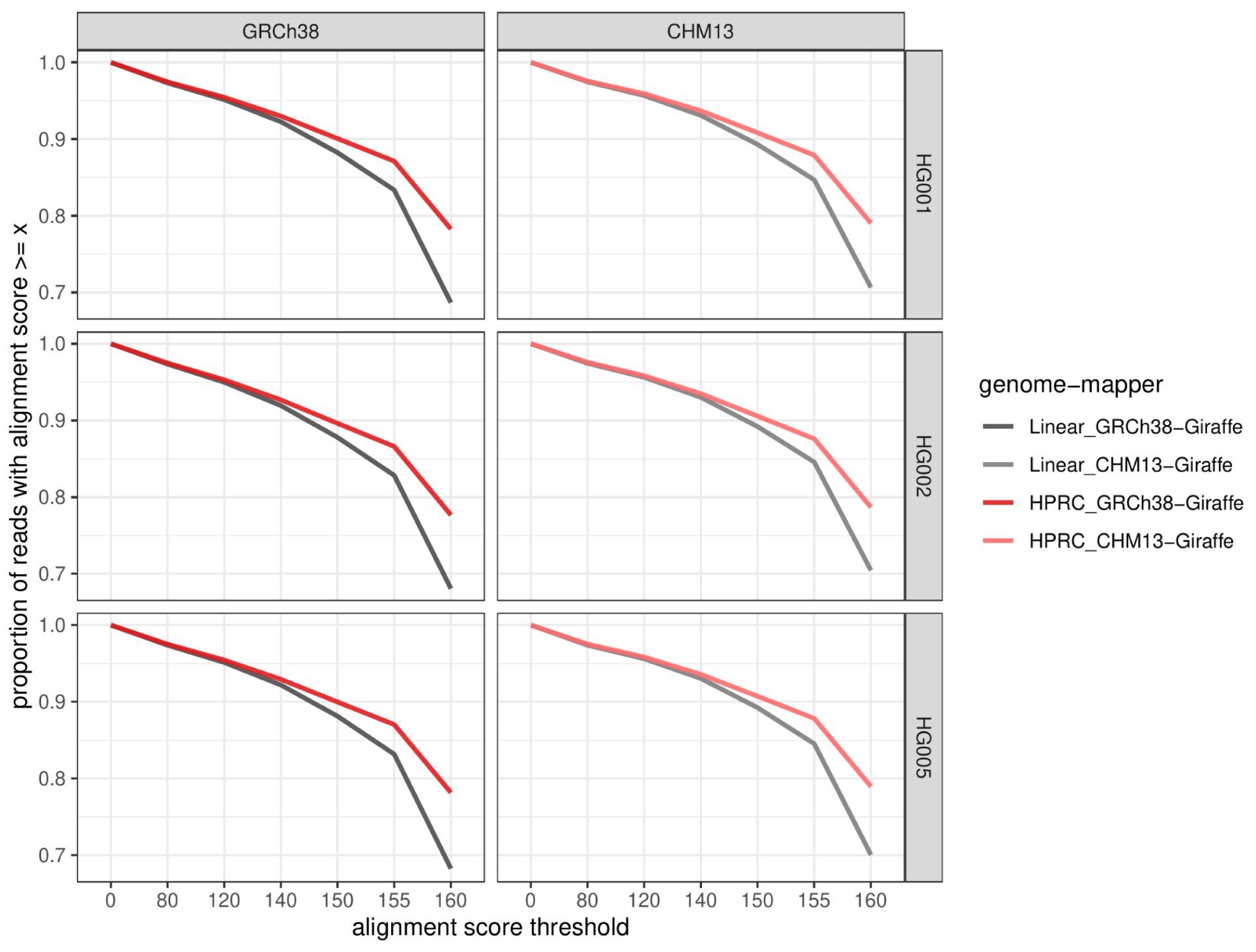
∼30x Illumina short-reads for three GIAB samples (horizontal panels) were mapped using two approaches: vg Giraffe on the linear pangenomes with just the reference genome (greys), and vg Giraffe on the HPRC pangenome (reds). The left panels compare GRCh38-referenced pangenomes, the right panels compare CHM13-referenced pangenomes. The curves show the proportion of reads (y-axis) with an alignment score greater or equal to the threshold defined by the x-axis.

**Supplementary Figure 6:**
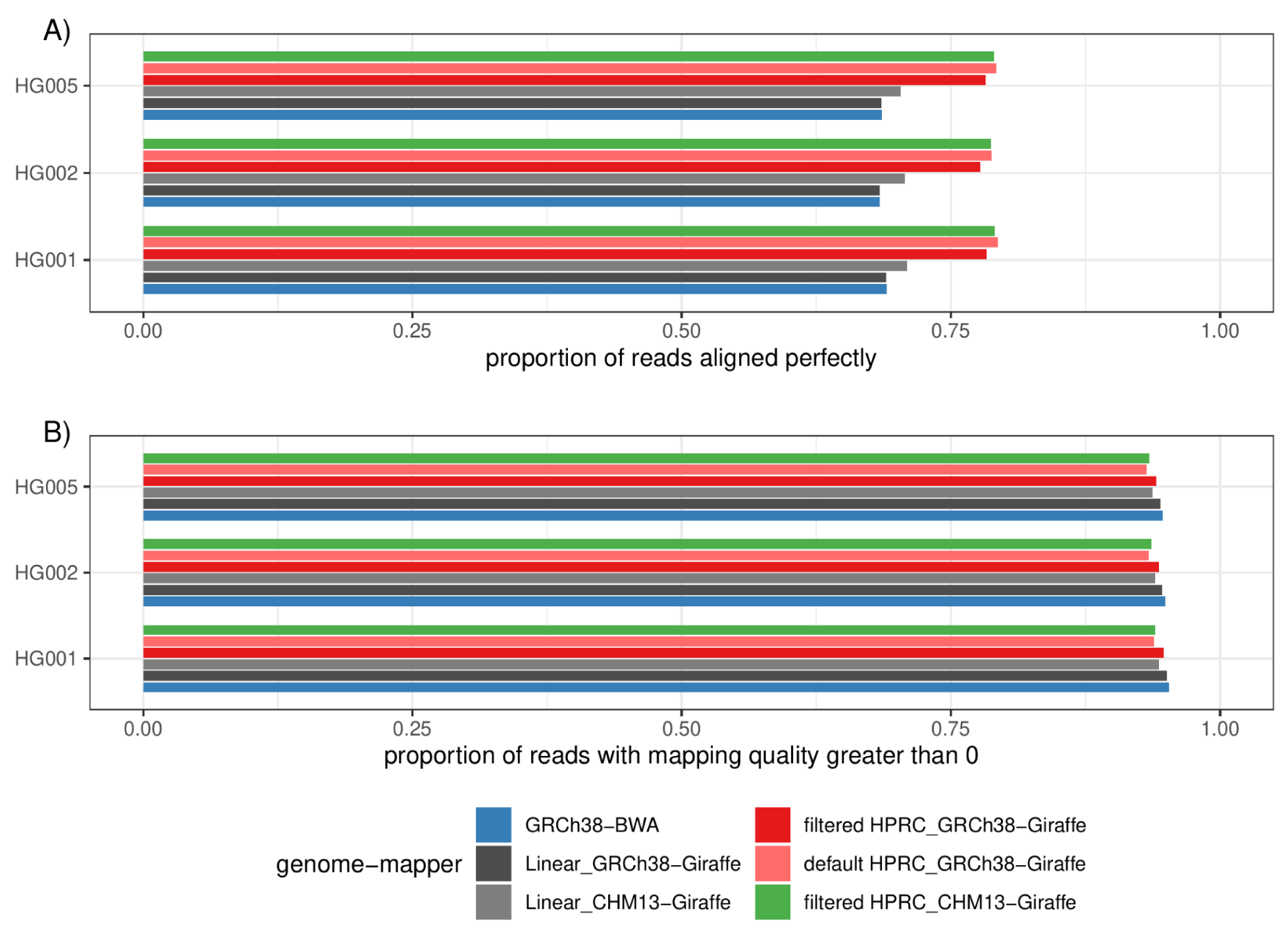
∼30x Illumina short-reads for three GIAB samples were mapped using three approaches: BWAMEM on GRCh38 (blue), vg Giraffe on the linear pangenomes with GRCh38 or CHM13 (grey), vg Giraffe on the GRCh38-referenced or CHM13-referenced HPRC pangenomes (red and green). The darker redbar corresponds to the default GRCh38-based HPRC pangenome, while the lighter redto the frequency-filtered pangenome used in practice for read mapping and variant calling. A) Proportion of the reads aligning perfectly to the (pan-)genome for each sample (y-axis). B) Proportion of reads with a mapping quality greater than 0.

**Supplementary Figure 7:**
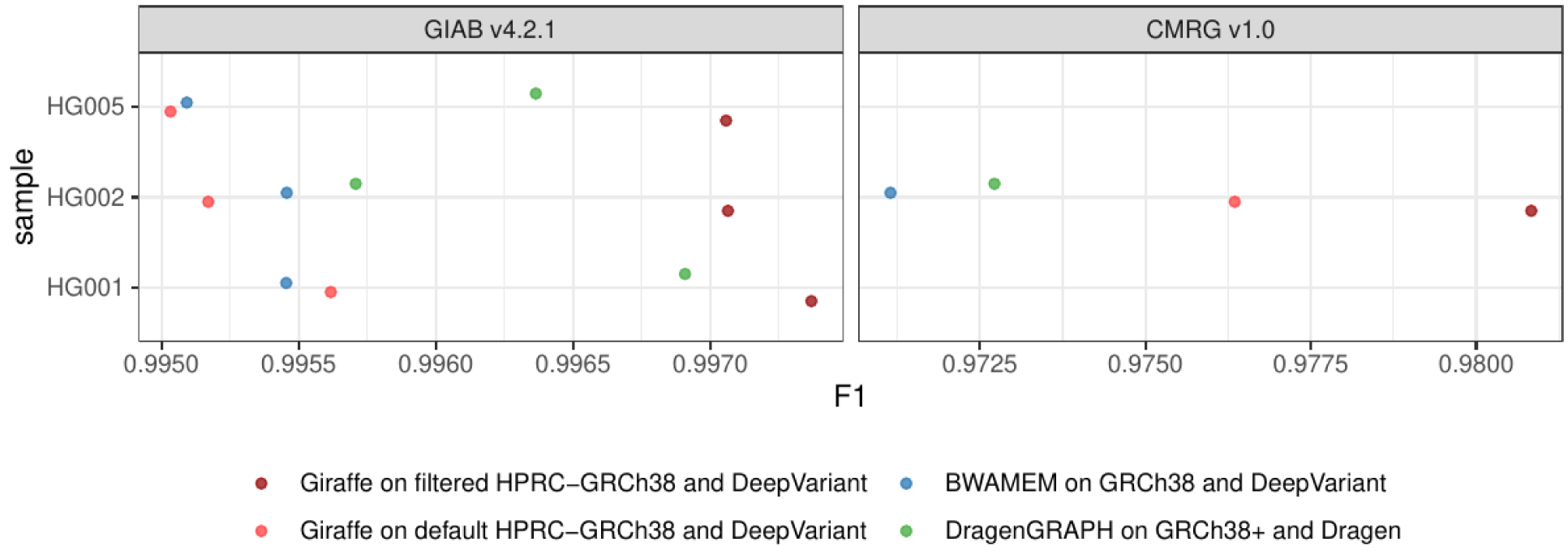
Evaluation of calls made on both the default pangenome (light red) and the frequency-filtered pangenome (dark red). The results when aligning reads with BWAMEM (blue) or using the Dragen pipeline (green) are also shown. The F1 score is shown on the x-axis across samples from the Genome in a Bottle (y-axis). Left: Genome in a Bottle v4.2.2 truth set. Right: Challenging Medically Relevant Genes v1.0 truth set.

**Supplementary Figure 8:**
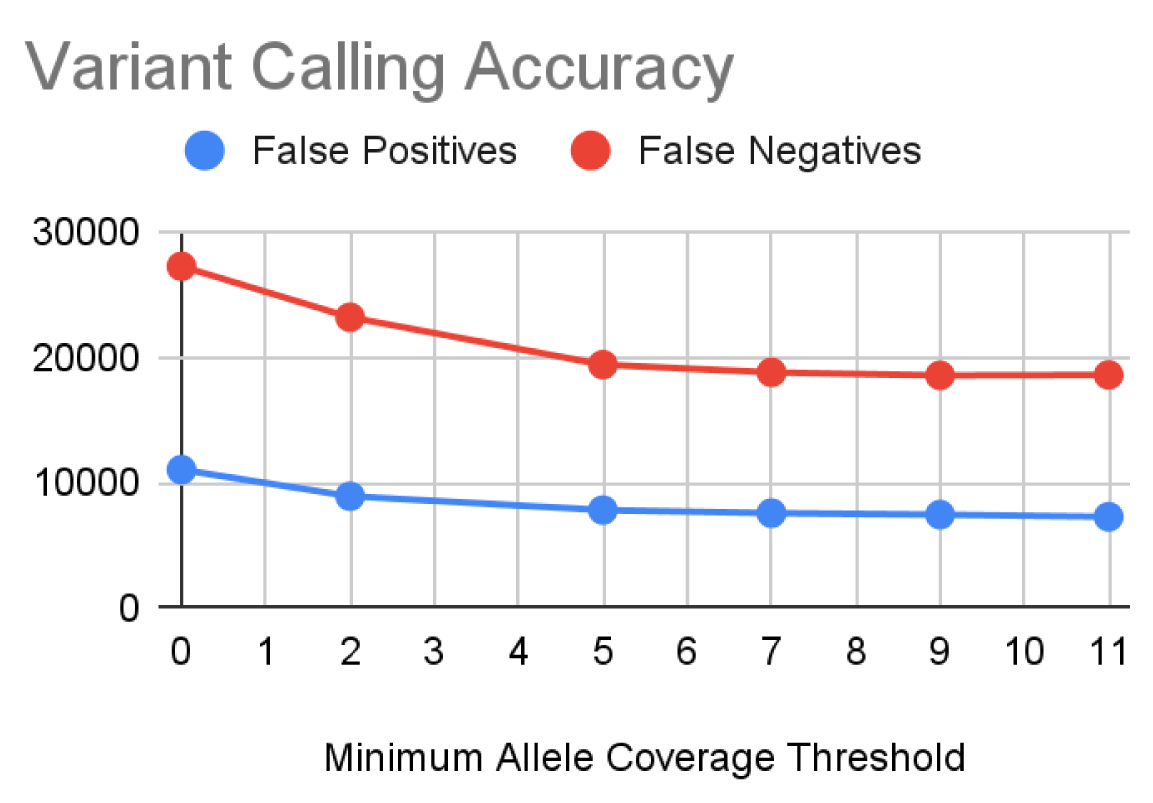
Variant calling false positives and false negatives on 30X Genome in a Bottle v4.2.1 Illumina reads for HG003 for the CHM13-based pangenome as a function of the allele frequency filtering threshold used. The 0 column is the unfiltered graph and the 9 column is the 10% (9/10) filter used for all other short-read mapping experiments. The accuracy was measured using rtg vcfeval v3.91^55^ on the evaluation regions provided by GIAB for this sample. The truth set has 3,831,915 calls total.

**Supplementary Figure 9:**
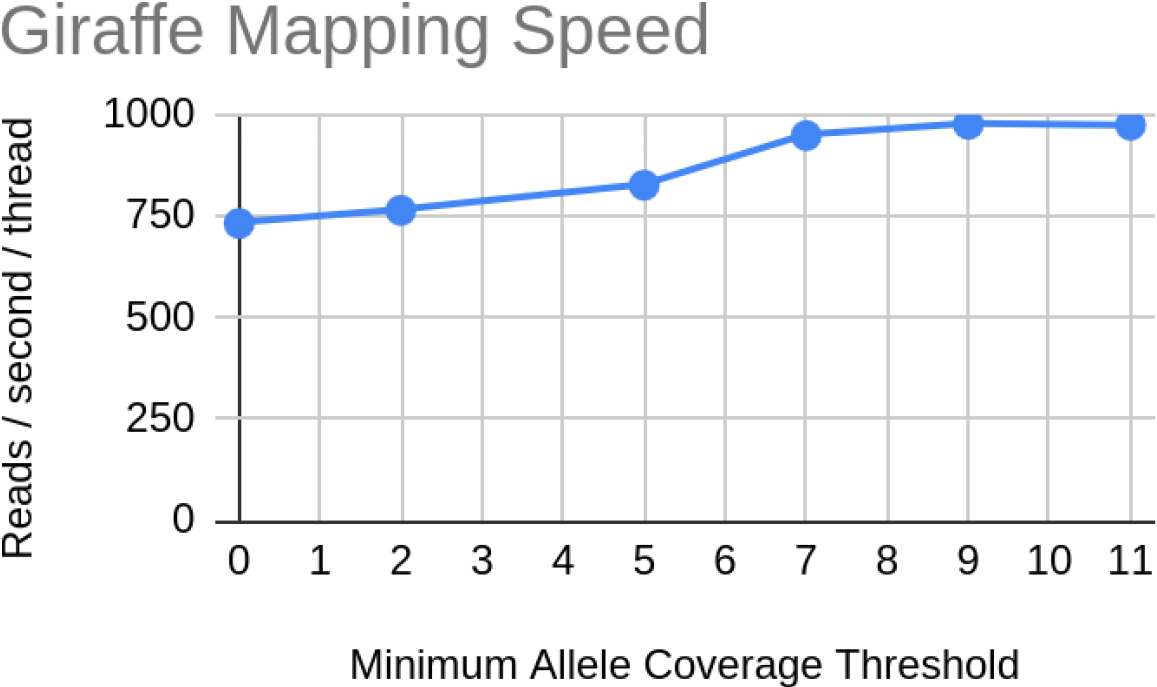
Short read mapping speed as a function of the allele frequency filtering threshold used, as reported by vg giraffe. The 0 column is the unfiltered graph and the 9 column is the 10% (9/10) filter used for all other short-read mapping experiments.

**Supplementary Figure 10:**
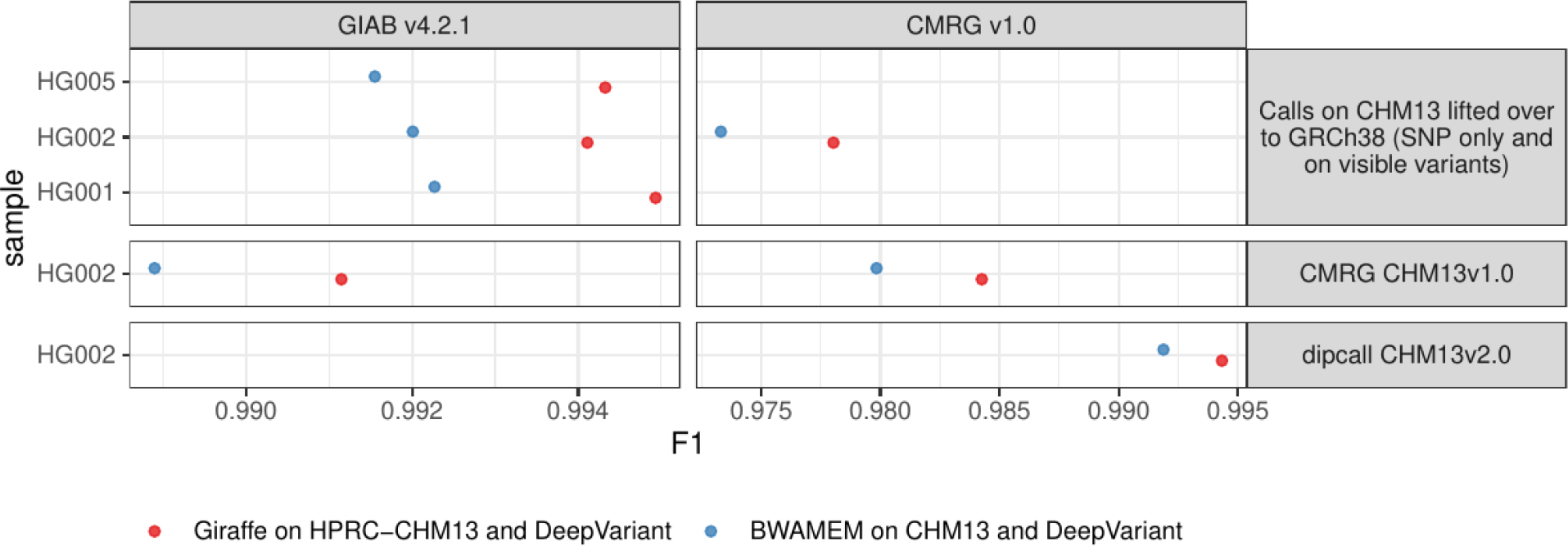
Evaluation of calls made on CHM13: aligning reads with BWAMEM (blue), or to the CHM13-based HPRC pangenome and projecting them to CHM13 (red). The F1 score is shown on the x-axis across samples from the Genome in a Bottle (y-axis). Left: Genome in a Bottle v4.2.2 truth set. Right: Challenging Medically Relevant Genes v1.0 truth set. Three approaches are shown as horizontal panels. Top: variants called on CHM13 were lifted over to be evaluated against the GRCh38 truth sets. Only SNPs and variant that are visible (not homozygous for the reference allele) on both reference genomes were used. Middle: the CMRG truth set for CHM13 v1.0 was lifted to CHM13 v2.0. The whole genome evaluation (left) was limited to the GIAB v4.2.1 confident regions lifted from GRCh38 to CHM13. Bottom: Preliminary draft truth set for CHM13 v2.0 based on HiFi assemblies analyzed with dipcall.

**Supplementary Figure 11:**
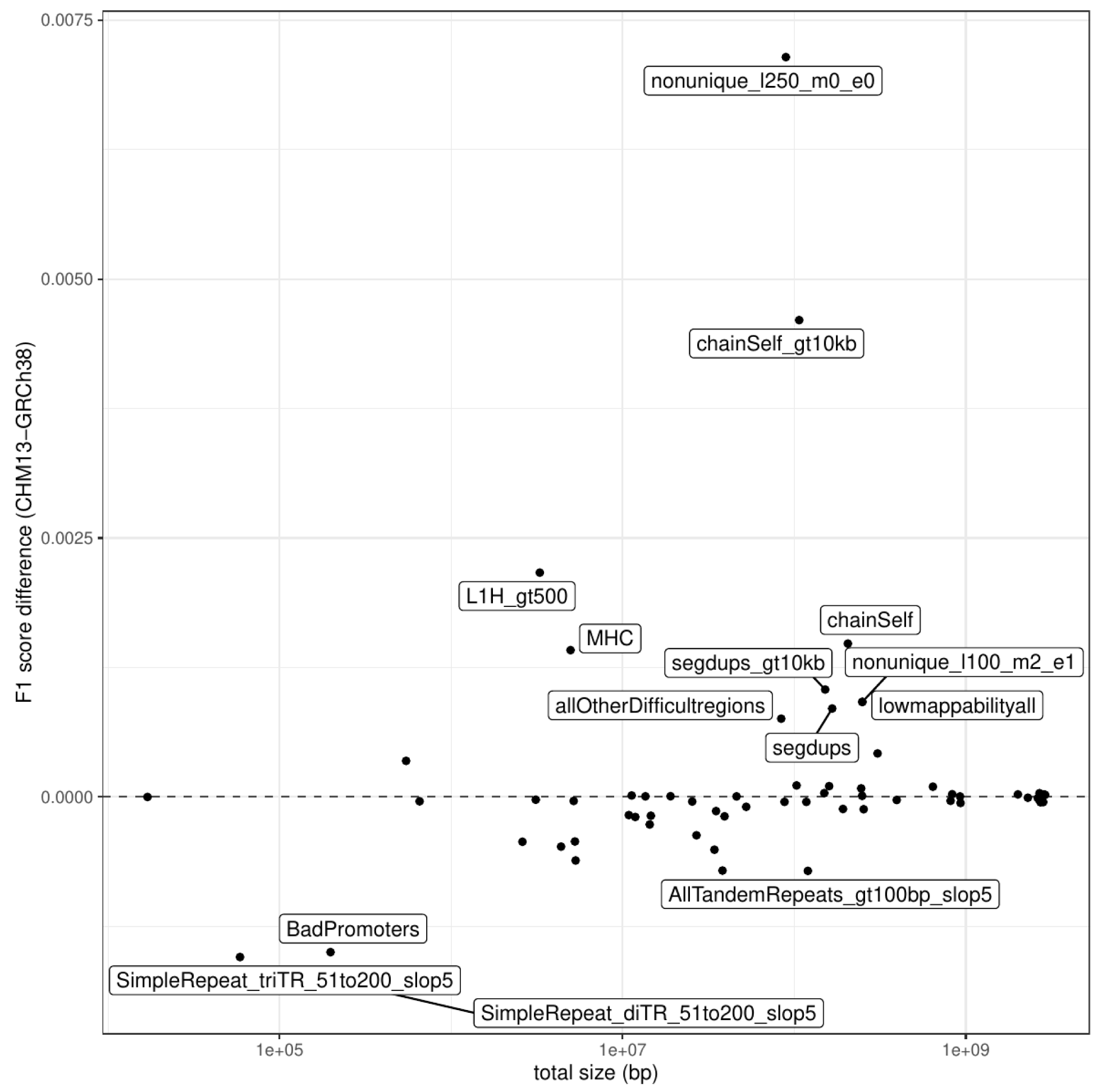
Difference between the F1 score obtained when using the CHM13-based pangenome compared to the GRCh38-based pangenome (y-axis), stratified by region sets from the GIAB (points). The total amount of sequence that represents each region set is shown on the x-axis. The top 10 most regions with the largest differences are labeled.

**Supplementary Figure 12:**
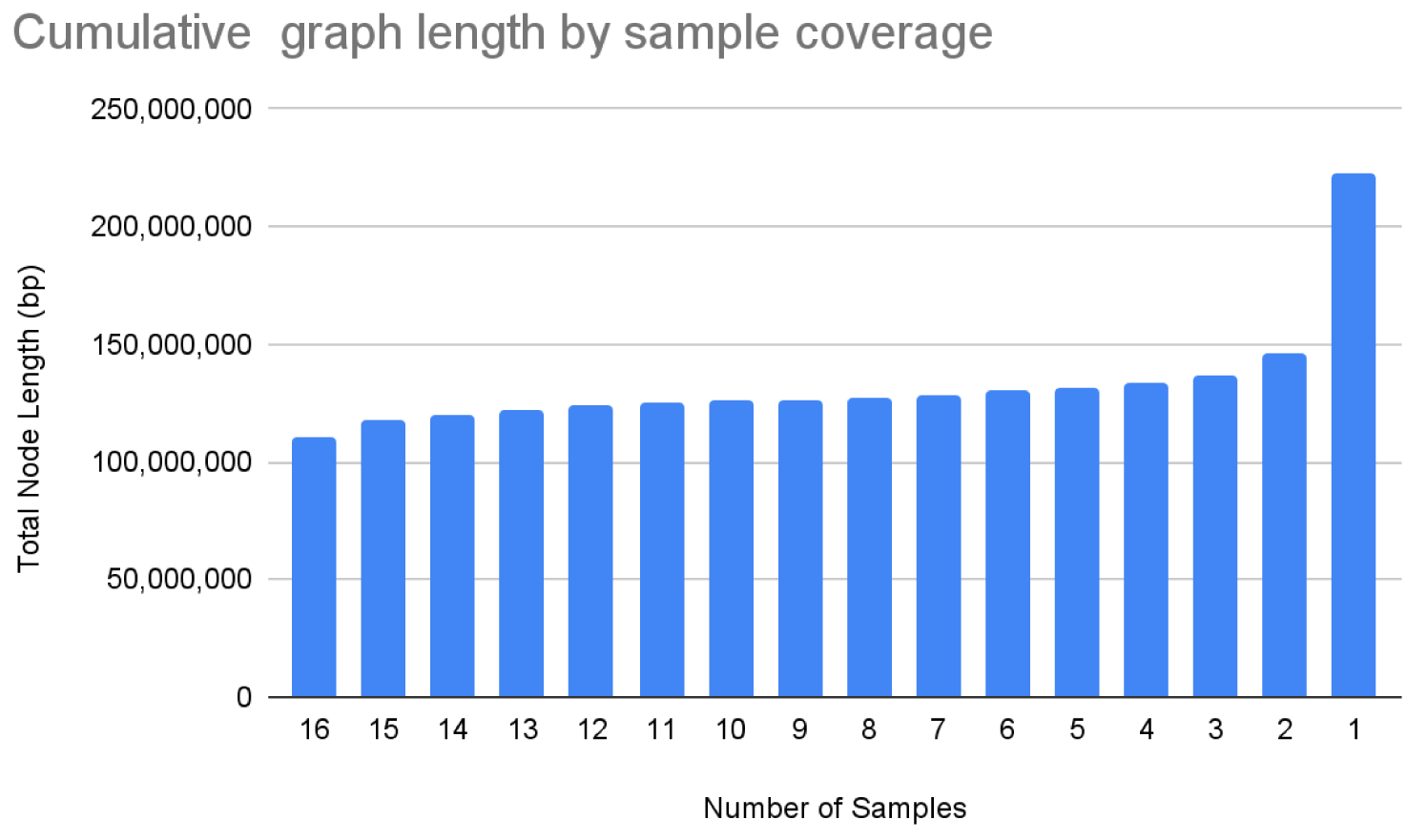
The amount of sequence in the *D. melanogaster* graph by the minimum number of haplotypes that contain it.

**Supplementary Figure 13:**
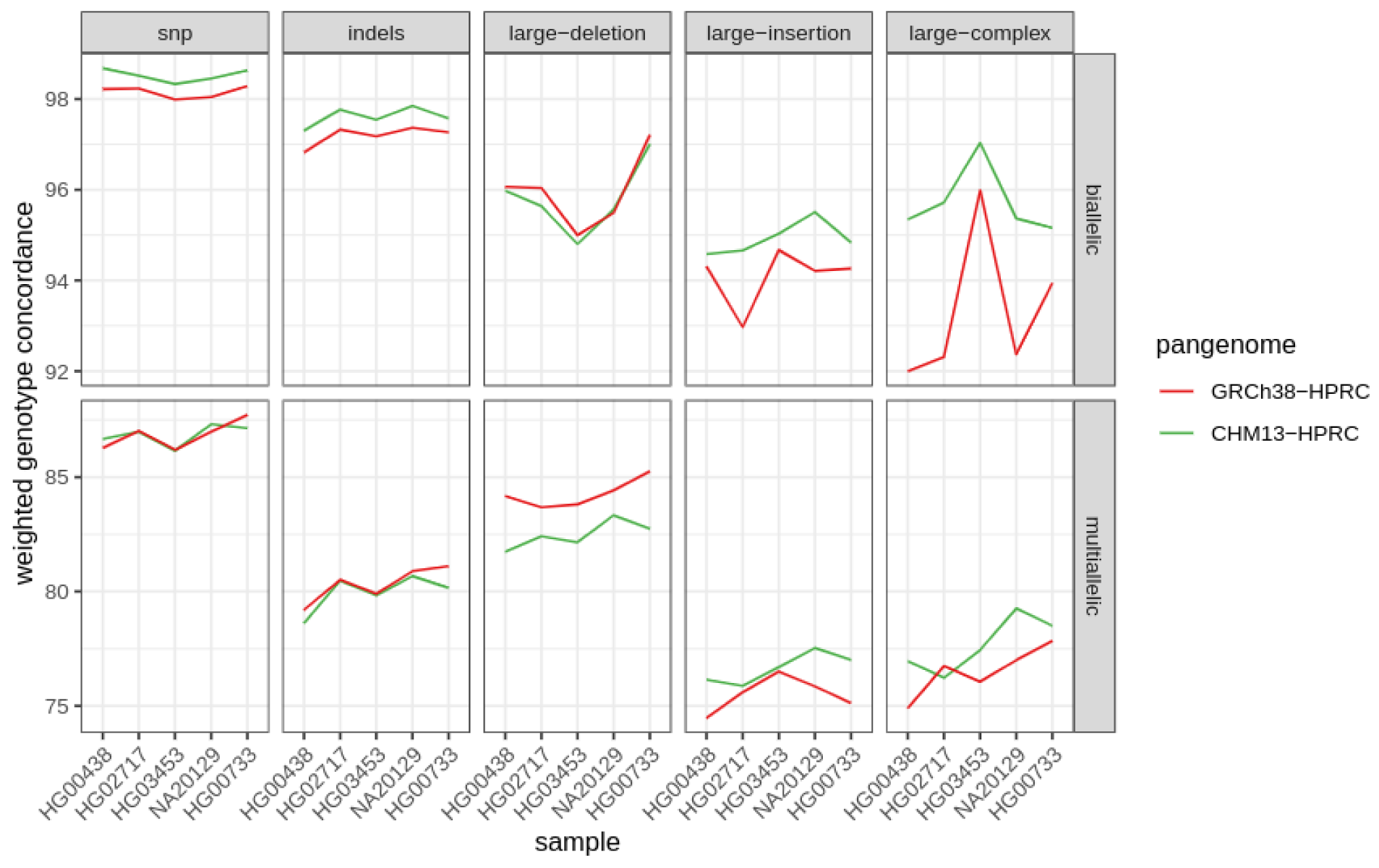
Leave-one-out PanGenie experiments comparing the GRCh38 and CHM13-based HPRC graphs across the same five samples. Each of the samples was, in turn, removed along with all its private variants from the input VCF to PanGenie then genotyped from short reads. The Weighted Genotype Concordance ^30^ was computed between the computed and original genotypes for each sample, excluding variants that were private (and therefore could not be re-genotyped). This is thus a comparison of the haplotypes as computed by Pangenie with those from the original assemblies within the context of the pangenome graph. Measurements are separated by variation category, as well as between bi-allelic and multiallelic sites in the graph.

**Supplementary Figure 14:**
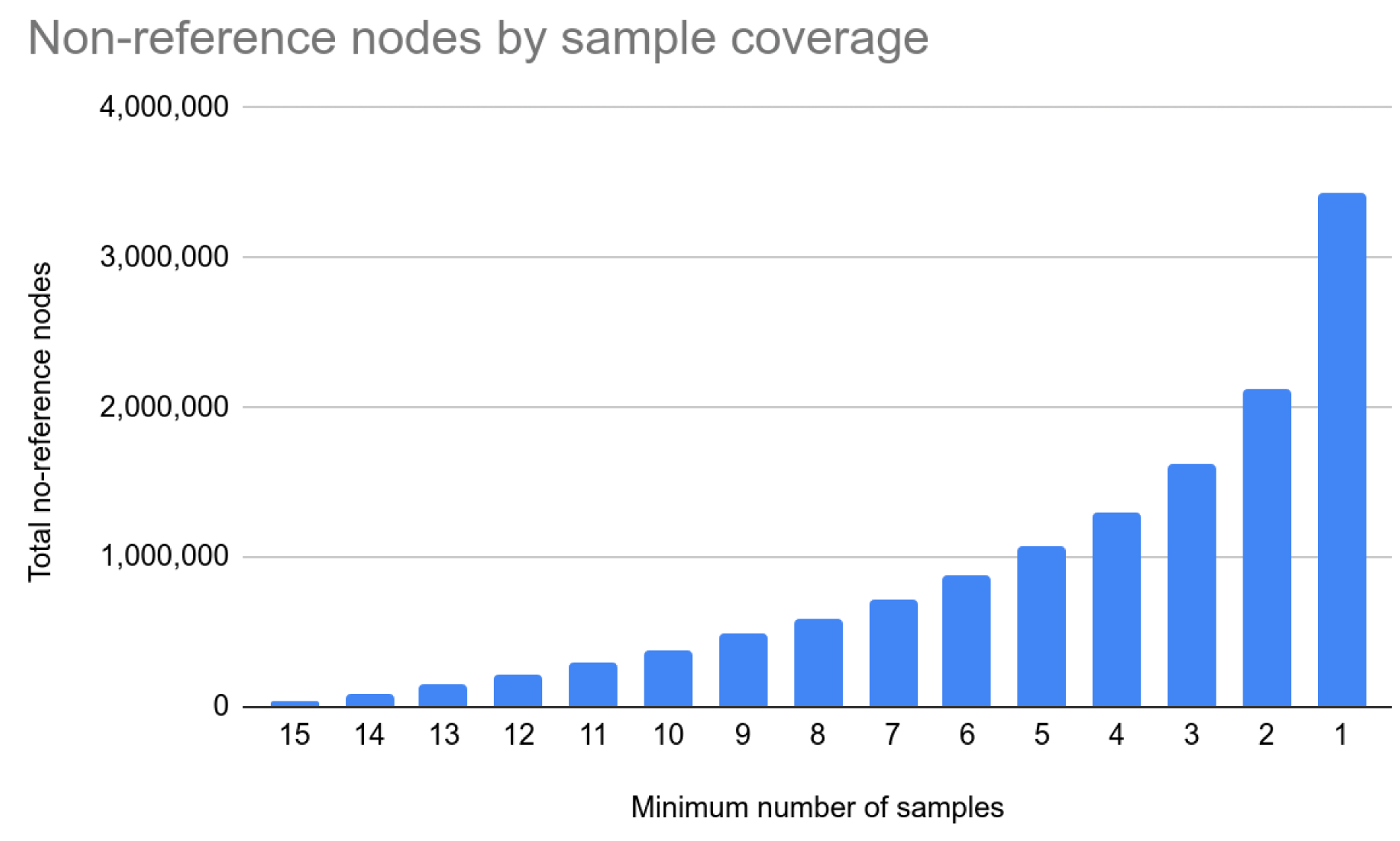
The number of nodes not present in dm6 covered by at least the given number of samples.

**Supplementary Figure 15:**
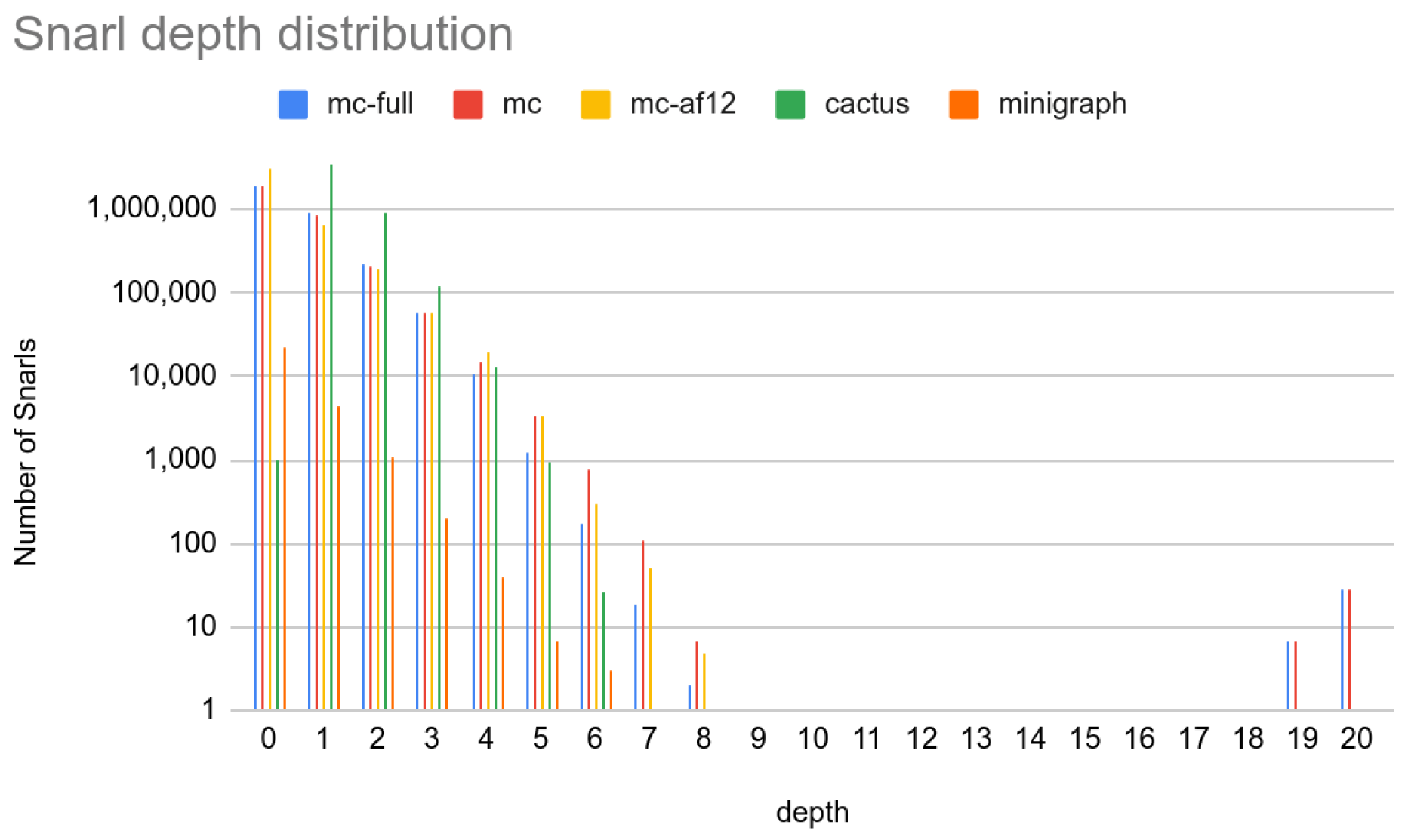
Snarl depth distribution.

**Supplementary Figure 16:**
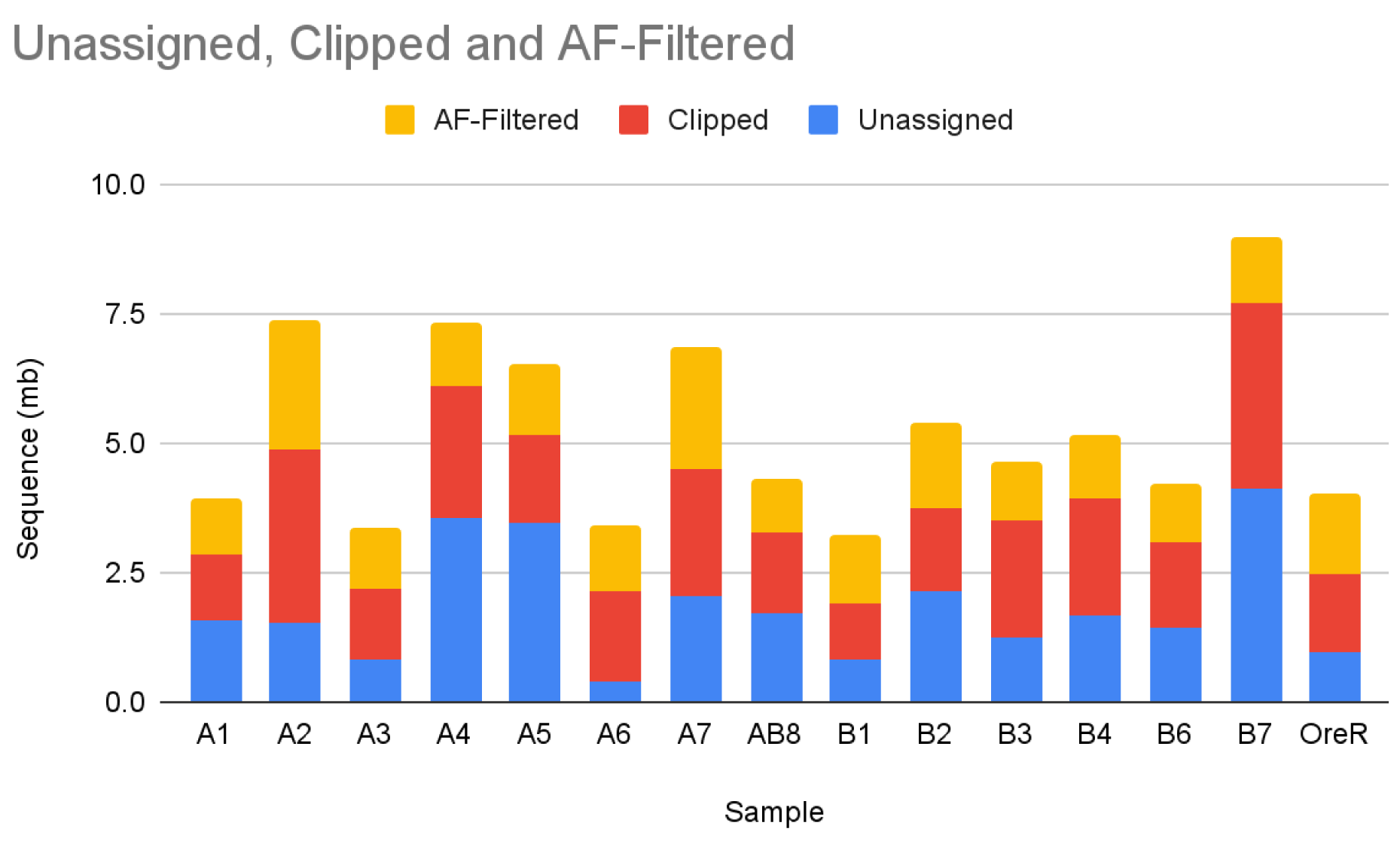
Sequence excluded from the *D. melanogaster* pangenome.

**Supplementary Figure 17:**
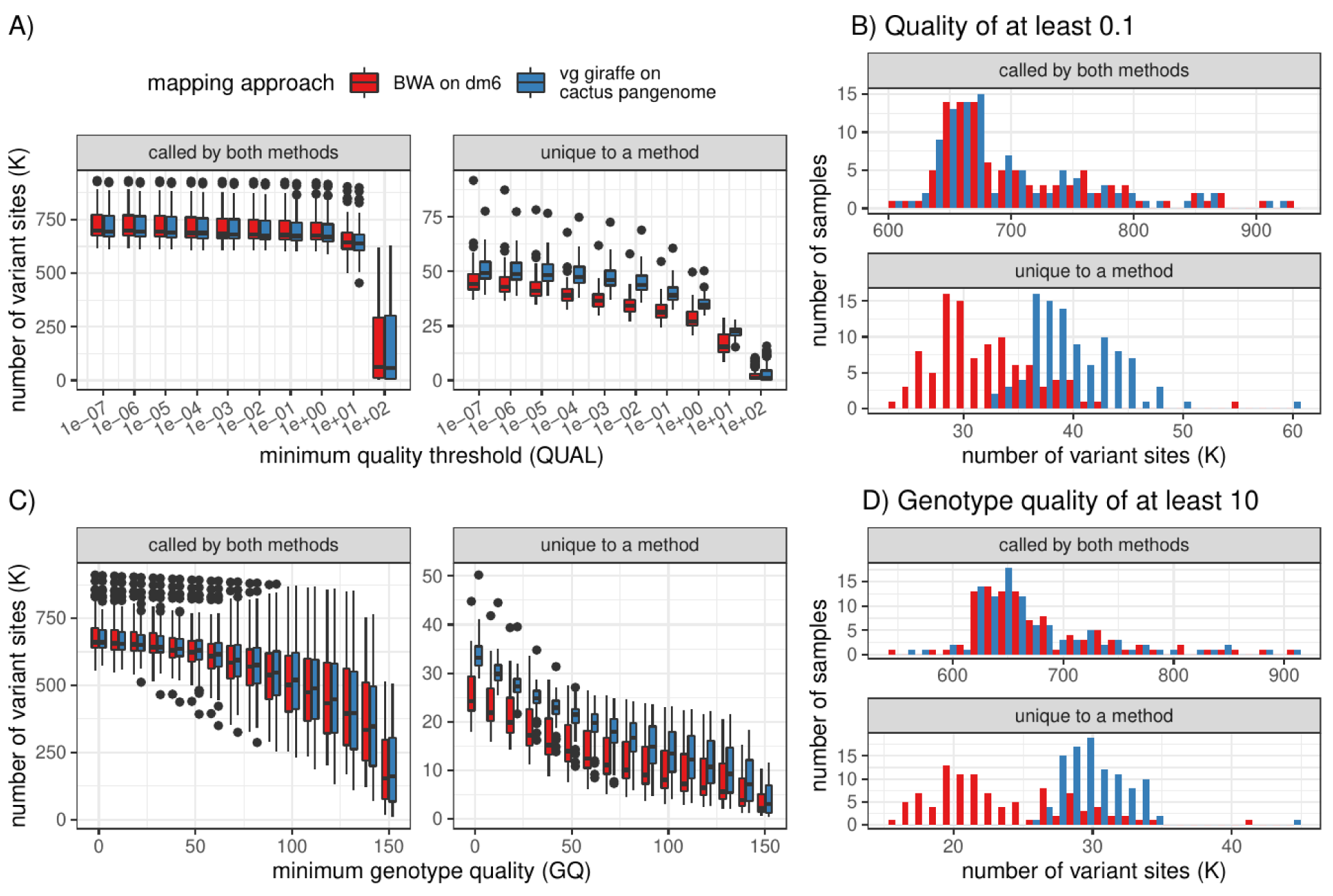
Number of variant sites with an alternate allele called in each of the 100 samples with FreeBayes. Two mapping approaches are compared: short-reads mapped to dm6 using BWA-MEM (red); short-reads mapped to the pangenome using vg Giraffe (blue). The variant sites were split into sites found by both approaches and sites found only by one. **A)** Distribution of the number of variant sites for different minimum quality (QUAL field) (x-axis). The boxplots show the median (center line), upper and lower quartiles (box limits), up to 1.5x interquartile range (whiskers), and outliers (points). **B)** Only variant sites with a quality of at least 0.1 were counted. This corresponds to x=0.1 in A). **C)** Distribution of the number of variant sites for different minimum genotype quality (GQ field) (x-axis). **D)** Only variant sites with a genotype quality of at least 10 were counted. This corresponds to x=10 in C).

**Supplementary Figure 18:**
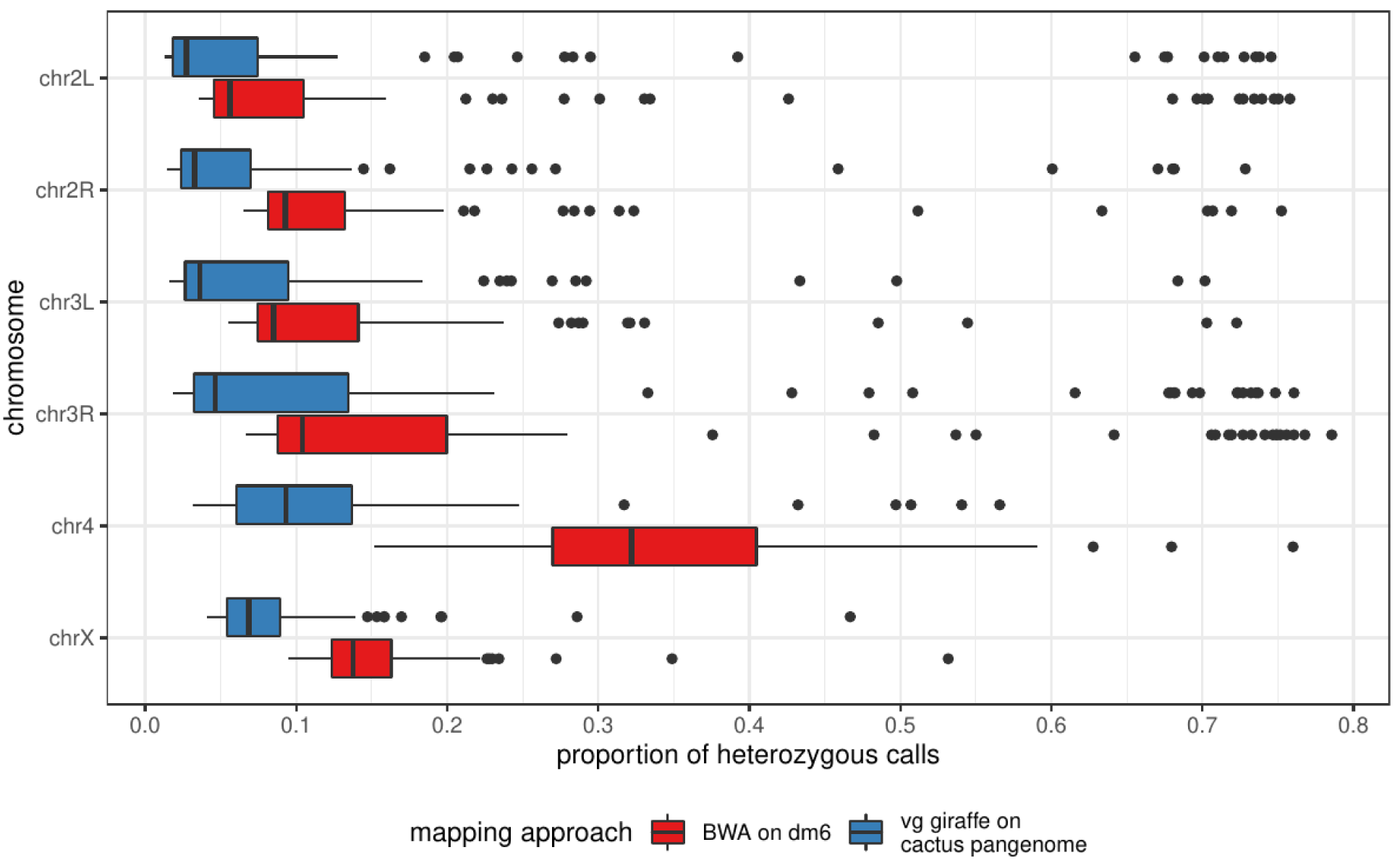
Proportion of heterozygous small variants called by FreeBayes in each of the 100 fly samples (point). Reads were either aligned to the pangenome and projected to dm6 (blue), or mapped to dm6 with BWA-MEM (red). Due to the inbreeding of these lines, we expect low heterozygosity. The boxplots show the median (center line), upper and lower quartiles (box limits), up to 1.5x interquartile range (whiskers), and outliers (points).

**Supplementary Figure 19:**
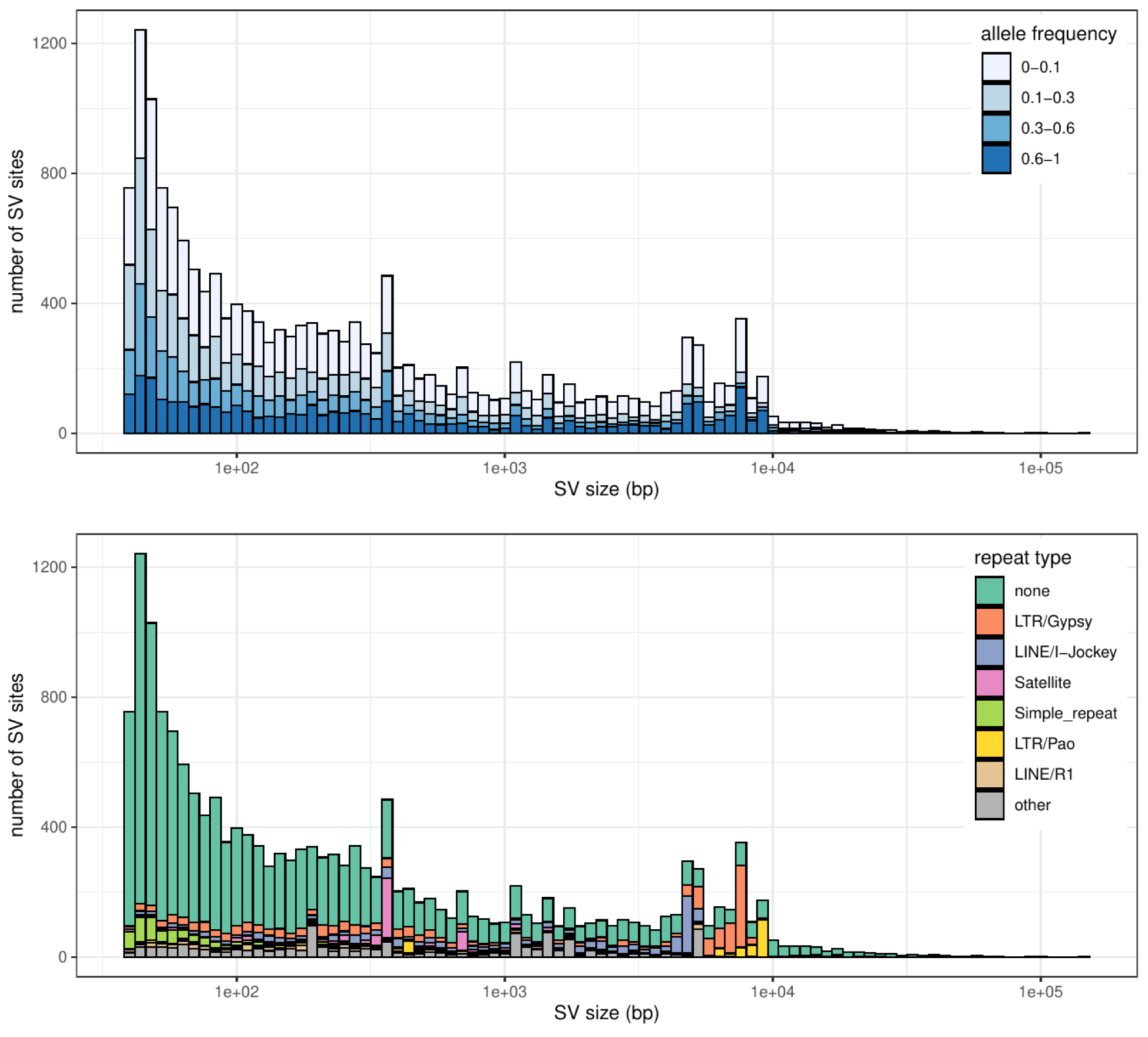
Distribution of the size of the SVs genotyped across 100 fly samples. The x-axis is log-scaled. **Top:** The SVs are colored by their allele frequencies. **Bottom:** The SVs are colored by the repeat class as annotated by Repeat Masker ^56^

**Supplementary Table 1:** HPRC graph sizes. The total node length is the sum of the lengths of all nodes in the graph. The total “non-ref” node length is the sum of the lengths of all nodes that are not present in any reference path, i.e. excluding CHM13 paths from the CHM13-based graph and GRCh38 paths from the GRCh38-based graph. The total path length is the sum of all paths in the graph, which will correspond exactly to the total length of all contigs input into the construction procedure, minus any sequence clipped out or unassignable to a chromosome.

| Graph | Nodes | Edges | Total Node Length | Total Non-ref Node Length | Total Path Length |
| --- | --- | --- | --- | --- | --- |
| SV Graph (GRCh38) | 424,643 | 637,628 | 3,239,764,787 | 140,014,069 | 3,239,764,787 |
| SV Graph (CHM13) | 493,631 | 738,529 | 3,365,688,482 | 253,629,026 | 3,365,688,482 |
| GRCh38-based Pangenome | 81,751,614 | 113,258,931 | 3,287,932,785 | 188,182,067 | 254,821,009,311 |
| GRCh38-based Filtered Pangenome (AF≥10%) | 59,960,908 | 72,408,601 | 3,153,443,019 | 53,692,301 | 254,415,272,646 |
| CHM13-based Pangenome | 85,591,995 | 118,409,526 | 3,324,657,754 | 212,598,298 | 257,143,252,360 |
| CHM13-based Filtered Pangenome (AF≥10%) | 62,335,399 | 75,270,997 | 3,166,744,316 | 54,684,860 | 256,673,009,341 |

**Supplementary Table 2:** *D. Melegonaster* graph sizes. The total node length is the sum of the lengths of all nodes in the graph. The total “non-ref” node length is the sum of the lengths of all nodes that are not present in any dm6 path. The total path length is the sum of all paths in the graph, which will correspond exactly to the total length of all contigs input into the construction procedure, minus any sequence clipped out or unassignable to a chromosome.

| Graph | Nodes | Edges | Total Node Length | Total Non-ref Node Length | Total Path Length |
| --- | --- | --- | --- | --- | --- |
| SV Graph | 80,853 | 112,742 | 214,547,800 | 71,326,590 | 214,547,800 |
| Unclipped Pangenome | 9,042,502 | 12,364,039 | 251,857,504 | 251,857,504 | 2,182,961,082 |
| Pangenome | 8,978,195 | 12,276,452 | 223,071,144 | 223,071,144 | 2,152,888,069 |
| Filtered Pangenome (AF12.5%) | 7,686,219 | 9,788,690 | 202,497,872 | 202,497,872 | 2,131,677,729 |
| Progressive Cactus Graph | 12,974,720 | 17,684,675 | 470,148,493 | 470,148,493 | 2,216,588,031 |

**Supplementary Table 3:** Number of variants in graphs genotyped by PanGenie. These statistics are obtained from the VCF after preprocessing by PanGenie. Indels are small evnets

| Variant type | HPRC-CHM13 |  |  | HPRC-GRCh38 |  |  | HGSVC-GRCh38 |  |  |
| --- | --- | --- | --- | --- | --- | --- | --- | --- | --- |
|  | # alleles in biallelic regions | # alleles in multiallelic regions | # alleles total | # alleles in biallelic regions | # alleles in multiallelic regions | # alleles total | # alleles in biallelic regions | # alleles in multiallelic regions | # alleles total |
| SNPs | 18,489,918 | 1,997,181 | **20,487,099** | 18,392,518 | 1,801,599 | **20,194,117** | 15,588,809 | 432,076 | **16,020,885** |
| Indels | 2,248,578 | 4,731,607 | **6,980,185** | 2,250,931 | 4,597,184 | **6,848,115** | 950,970 | 232,447 | **1,183,417** |
| SV-DEL | 4,270 | 66,440 | **70,710** | 4,232 | 52,969 | **57,201** | 6,685 | 29,174 | **35,859** |
| SV-INS | 12,384 | 187,443 | **199,827** | 12,192 | 242,420 | **254,612** | 37,340 | 23,078 | **60,418** |
| SV-Other | 1,382 | 90,766 | **92,148** | 1,296 | 100,700 | **101,996** | - | - | - |

**Supplementary Table 4:**
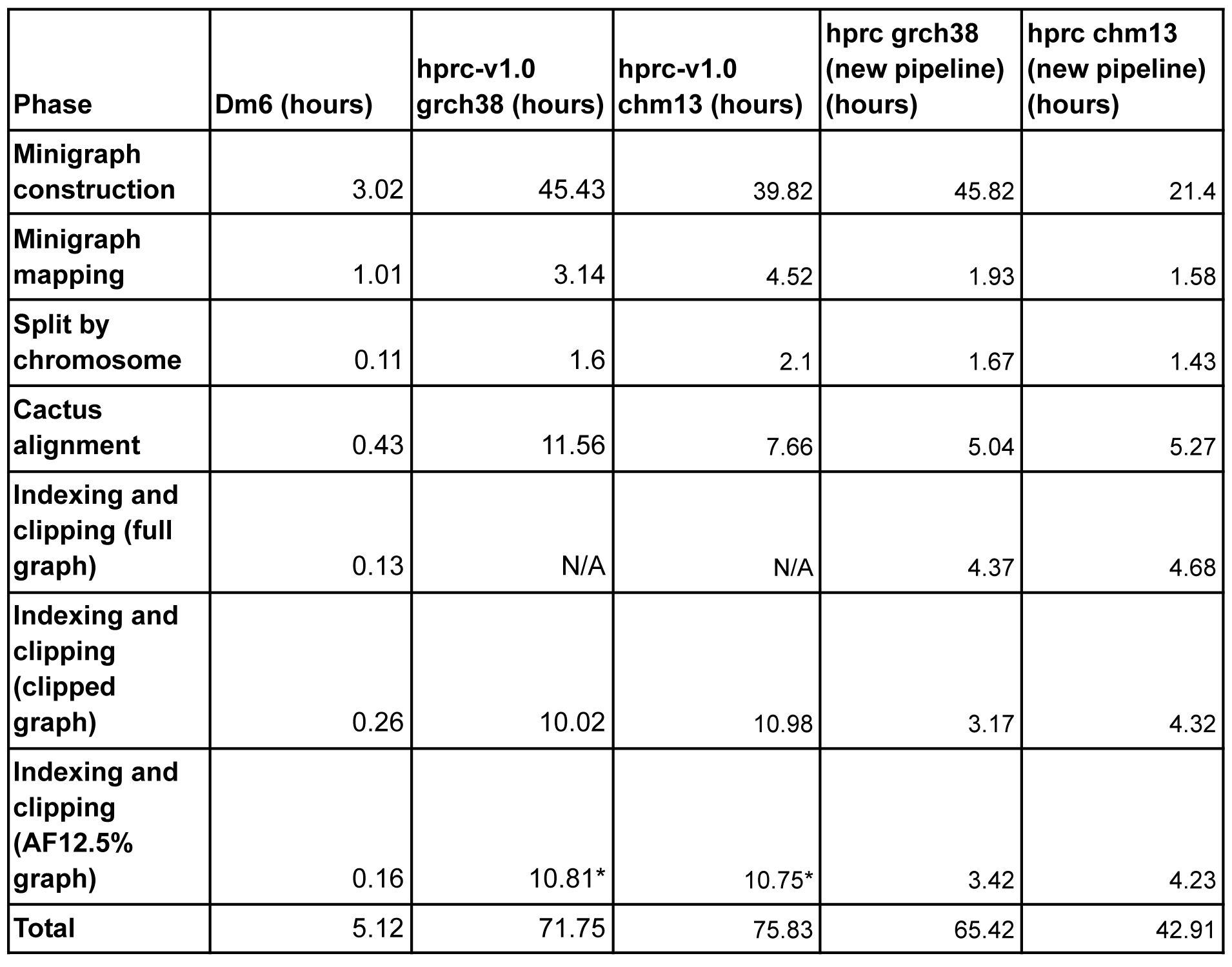
Minigraph-Cactus running times (wall-times). The “new pipeline” columns refer to graphs made using the method described here which does not rely on dna-brnn for clipping. The dm6 graphs were made using up to 32 cores and 16Gb RAM. The HPRC graphs were made on an AWS cluster using up to 25 32 core 256Gb RAM machines, except for the indexing stages which were done on up to 2 64 core 512Gb RAM machines. The disk usage of each step is bounded by the total size of the input and output (plus uncompressed versions of the same if they are gzipped). *\* These values were not kept in the logs and were estimated using the ratios in the neighboring columns (ex 10.81 = 3.42/3.17 * 10.02)*.

**Supplementary Table 5:** Progressive Cactus running times (wall times) using single 32-core machine with up to 64Gb RAM.

| Phase | Dm6 (hours) |
| --- | --- |
| Lastz repeatmasking | 0.38 |
| All-to-all lastz alignment | 17.97 |
| Cactus alignment | 0.83 |
| Total | 19.18 |

**Supplementary Table 6:** DGRP sequencing data used for *D. Melanogaster* mapping and variant calling experiments

| DGRP Line | Sequencing Technology | Freeze | Mapped Coverage | Raw Read Length:Read Number | NCBI SRA | NCBI SRR |
| --- | --- | --- | --- | --- | --- | --- |
| DGRP_21 | Illumina | F1 | 15.8 | 95bp:37046984 | SRX021040 | SRR834526 |
| DGRP_31 | Illumina | F2 | 49.2 | 125bp:76894692 | SRX155996 | SRR834509 |
| DGRP_32 | Illumina | F2 | 56.2 | 125bp:88154526 | SRX155997 | SRR834512 |
| DGRP_38 | Illumina | F1 | 28.0 | 95bp:56154204 | SRX025317 | SRR834541 |
| DGRP_40 | Illumina | F1 | 33.3 | 95bp:69063428 | SRX021235 | SRR835025 |
| DGRP_42 | Illumina | F1 | 20.2 | 95bp:37186556 | SRX021255 | SRR835027 |
| DGRP_48 | Illumina | F2 | 32.7 | 125bp:58419132 | SRX155989 | SRR835034 |
| DGRP_49 | Illumina | F1 | 15.2 | 75bp:37870818 | SRX021267 | SRR835037 |
| DGRP_57 | Illumina | F1 | 32.6 | 100bp:64966990 | SRX021296 | SRR933581 |
| DGRP_75 | Illumina | F1 | 18.5 | 110bp:38161744 | SRX021384 | SRR835087 |
| DGRP_83 | Illumina | F1 | 16.3 | 75bp:41070470 | SRX023456 | SRR835058 |
| DGRP_100 | Illumina | F2 | 52.3 | 125bp:87340978 | SRX156026 | SRR833244 |
| DGRP_138 | Illumina | F1 | 30.1 | 100bp:61689820 | SRX021008 | SRR932121 |
| DGRP_142 | Illumina | F1 | 19.7 | 110bp:41167794 | SRX020759 | SRR834551 |
| DGRP_177 | Illumina | F1 | 24.6 | 95bp:49114764 | SRX021026 | SRR834547 |
| DGRP_181 | Illumina | F1 | 24.7 | 75bp:64093862 | SRX020912 | SRR933563 |
| DGRP_189 | Illumina | F2 | 37.8 | 125bp:63289120 | SRX155979 | SRR834523 |
| DGRP_223 | Illumina | F2 | 40.8 | 125bp:71152512 | SRX155994 | SRR834527 |
| DGRP_235 | Illumina | F1 | 18.4 | 95bp:38296004 | SRX021053 | SRR834531 |
| DGRP_318 | Illumina | F1 | 15.2 | 75bp:39068236 | SRX021082 | SRR834507 |
| DGRP_319 | Illumina | F2 | 37.6 | 125bp:70621686 | SRX155981 | SRR834508 |
| DGRP_320 | Illumina | F1 | 24.2 | 95bp:51875680 | SRX021063 | SRR834510 |
| DGRP_321 | Illumina | F1 | 33.5 | 95bp:67314152 | SRX021094 | SRR834511 |
| DGRP_332 | Illumina | F1 | 25.7 | 75bp:65583082 | SRX021095 | SRR933569 |
| DGRP_348 | Illumina | F2 | 48.3 | 125bp:78515972 | SRX156029 | SRR834514 |
| DGRP_352 | Illumina | F1 | 15.6 | 75bp:44982388 | SRX021101 | SRR834516 |
| DGRP_354 | Illumina | F2 | 57.2 | 101bp:106369344 | SRX156027 | SRR834517 |
| DGRP_355 | Illumina | F2 | 44.9 | 101bp:84541222 | SRX156028 | SRR834545 |
| DGRP_356 | Illumina | F1 | 15.5 | 75bp:42903612 | SRX023833 | SRR834537 |
| DGRP_359 | Illumina | F1 | 20.2 | 95bp:37271884 | SRX023424 | SRR834546 |
| DGRP_361 | Illumina | F2 | 40.6 | 125bp:68254340 | SRX155984 | SRR834553 |
| DGRP_370 | Illumina | F1 | 20.9 | 95bp:43793604 | SRX021104 | SRR834539 |
| DGRP_377 | Illumina | F1 | 21.8 | 95bp:43796182 | SRX023834 | SRR834543 |
| DGRP_381 | Illumina | F1 | 20.9 | 75bp:54335852 | SRX021112 | SRR933573 |
| DGRP_382 | Illumina | F2 | 41.1 | 125bp:73812254 | SRX156013 | SRR834552 |
| DGRP_383 | Illumina | F1 | 19.1 | 95bp:39897030 | SRX021113 | SRR834554 |
| DGRP_390 | Illumina | F2 | 26.2 | 125bp:42709922 | SRX156014 | SRR834519 |
| DGRP_392 | Illumina | F1 | 23.2 | 95bp:51156860 | SRX021157 | SRR834520 |
| DGRP_395 | Illumina | F2 | 47.1 | 101bp:87233368 | SRX156015 | SRR834521 |
| DGRP_397 | Illumina | F2 | 30.0 | 125bp:48910026 | SRX156017 | SRR834522 |
| DGRP_405 | Illumina | F1 | 22.9 | 95bp:50080536 | SRX021242 | SRR835023 |
| DGRP_406 | Illumina | F1 | 25.0 | 95bp:51821248 | SRX021254 | SRR835024 |
| DGRP_426 | Illumina | F1 | 21.1 | 95bp:43746634 | SRX021245 | SRR835026 |
| DGRP_427 | Illumina | F1 | 16.3 | 45bp:64106936 | SRX006155 | SRR933577 |
| DGRP_439 | Illumina | F1 | 20.4 | 95bp:44762436 | SRX021244 | SRR835028 |
| DGRP_440 | Illumina | F1 | 17.2 | 95bp:43161850 | SRX021246 | SRR835029 |
| DGRP_441 | Illumina | F1 | 18.7 | 95bp:42278010 | SRX023835 | SRR835030 |
| DGRP_443 | Illumina | F1 | 28.5 | 95bp:57567568 | SRX021260 | SRR835031 |
| DGRP_461 | Illumina | F1 | 21.9 | 95bp:49324528 | SRX021262 | SRR835033 |
| DGRP_491 | Illumina | F1 | 15.1 | 75bp:40944392 | SRX021268 | SRR835035 |
| DGRP_492 | Illumina | F1 | 22.1 | 95bp:44580310 | SRX021270 | SRR835036 |
| DGRP_502 | Illumina | F1 | 21.7 | 95bp:44336646 | SRX021271 | SRR835038 |
| DGRP_505 | Illumina | F2 | 43.7 | 125bp:71295212 | SRX156002 | SRR835039 |
| DGRP_508 | Illumina | F1 | 21.2 | 95bp:42338556 | SRX021272 | SRR835040 |
| DGRP_509 | Illumina | F1 | 15.3 | 75bp:38095912 | SRX021273 | SRR835041 |
| DGRP_513 | Illumina | F1 | 19.6 | 95bp:42640722 | SRX021282 | SRR835042 |
| DGRP_528 | Illumina | F2 | 36.2 | 125bp:57697778 | SRX155985 | SRR835043 |
| DGRP_530 | Illumina | F2 | 20.7 | 125bp:34726088 | SRX156031 | SRR835044 |
| DGRP_531 | Illumina | F1 | 17.9 | 95bp:41560152 | SRX021290 | SRR835045 |
| DGRP_535 | Illumina | F1 | 15.2 | 75bp:40234802 | SRX021293 | SRR835046 |
| DGRP_551 | Illumina | F2 | 21.4 | 125bp:35225968 | SRX156034 | SRR835047 |
| DGRP_555 | Illumina | F1 | 19.2 | 75bp:50103810 | SRX006159 | SRR933580 |
| DGRP_559 | Illumina | F2 | 24.2 | 125bp:36482062 | SRX156032 | SRR835048 |
| DGRP_566 | Illumina | F2 | 48.8 | 101bp:89414580 | SRX156033 | SRR835050 |
| DGRP_596 | Illumina | F2 | 41.1 | 101bp:73915046 | SRX156004 | SRR835096 |
| DGRP_627 | Illumina | F2 | 36.7 | 125bp:82297368 | SRX155988 | SRR835097 |
| DGRP_630 | Illumina | F2 | 21.7 | 125bp:36162916 | SRX156003 | SRR835098 |
| DGRP_634 | Illumina | F2 | 19.4 | 125bp:32632568 | SRX156018 | SRR835086 |
| DGRP_705 | Illumina | F1 | 16.7 | 75bp:47006608 | SRX006162 | SRR933585 |
| DGRP_707 | Illumina | F1 | 17.8 | 75bp:46657404 | SRX006163 | SRR933586 |
| DGRP_712 | Illumina | F1 | 16.3 | 75bp:44687868 | SRX006164 | SRR933587 |
| DGRP_727 | Illumina | F1 | 27.5 | 75bp:73781476 | SRX021382 | SRR933589 |
| DGRP_732 | Illumina | F1 | 16.3 | 75bp:42170344 | SRX006167 | SRR933591 |
| DGRP_737 | Illumina | F1 | 25.1 | 75bp:74740132 | SRX023451 | SRR933592 |
| DGRP_738 | Illumina | F1 | 27.1 | 75bp:75804508 | SRX021383 | SRR933593 |
| DGRP_757 | Illumina | F1 | 28.4 | 75bp:74326240 | SRX021385 | SRR933594 |
| DGRP_761 | Illumina | F1 | 15.2 | 75bp:40867250 | SRX021386 | SRR835088 |
| DGRP_776 | Illumina | F1 | 15.6 | 75bp:39890986 | SRX021387 | SRR835089 |
| DGRP_787 | Illumina | F1 | 15.4 | 75bp:39795416 | SRX021388 | SRR835091 |
| DGRP_790 | Illumina | F1 | 17.0 | 95bp:35620658 | SRX021389 | SRR835092 |
| DGRP_805 | Illumina | F1 | 16.1 | 75bp:43182102 | SRX021400 | SRR835095 |
| DGRP_810 | Illumina | F1 | 15.5 | 75bp:36972402 | SRX021418 | SRR835051 |
| DGRP_812 | Illumina | F1 | 16.1 | 75bp:38719004 | SRX021419 | SRR835052 |
| DGRP_819 | Illumina | F2 | 73.0 | 100bp:150745358 | SRX156006 | SRR835054 |
| DGRP_822 | Illumina | F1 | 17.7 | 110bp:41079524 | SRX021476 | SRR835055 |
| DGRP_837 | Illumina | F1 | 20.7 | 95bp:46411538 | SRX021479 | SRR933599 |
| DGRP_843 | Illumina | F2 | 42.3 | 125bp:68658714 | SRX156036 | SRR835059 |
| DGRP_849 | Illumina | F2 | 39.9 | 125bp:61687178 | SRX156035 | SRR835060 |
| DGRP_850 | Illumina | F2 | 43.6 | 125bp:69699750 | SRX155993 | SRR835061 |
| DGRP_855 | Illumina | F1 | 19.2 | 110bp:42348166 | SRX021563 | SRR835062 |
| DGRP_857 | Illumina | F1 | 20.8 | 110bp:42340250 | SRX021492 | SRR835063 |
| DGRP_882 | Illumina | F1 | 17.4 | 75bp:44722234 | SRX021496 | SRR835067 |
| DGRP_887 | Illumina | F1 | 19.5 | 95bp:43595728 | SRX021527 | SRR835069 |
| DGRP_890 | Illumina | F1 | 15.9 | 75bp:41954706 | SRX021499 | SRR835071 |
| DGRP_892 | Illumina | F1 | 20.5 | 95bp:45702226 | SRX023838 | SRR835072 |
| DGRP_894 | Illumina | F1 | 16.8 | 95bp:35128536 | SRX021528 | SRR835073 |
| DGRP_897 | Illumina | F1 | 27.0 | 75bp:70892788 | SRX023457 | SRR933601 |
| DGRP_907 | Illumina | F1 | 17.5 | 95bp:36385056 | SRX021500 | SRR835074 |
| DGRP_908 | Illumina | F1 | 19.9 | 95bp:39111536 | SRX021501 | SRR835075 |
| DGRP_913 | Illumina | F2 | 43.7 | 125bp:69250292 | SRX156024 | SRR835077 |

## Bibliography

1. Eizenga JM, Novak AM, Sibbesen JA, Heumos S, Ghaffaari A, Hickey G et al. Pangenome Graphs. Annu Rev Genomics Hum Genet 2020; 21: 139–162.

2. Miga KH, Wang T. The Need for a Human Pangenome Reference Sequence. Annu Rev Genomics Hum Genet 2021; 22: 81–102.

3. Garrison E, Sirén J, Novak AM, Hickey G, Eizenga JM, Dawson ET et al. Variation graph toolkit improves read mapping by representing genetic variation in the reference. Nat Biotechnol 2018; 36: 875–879.

4. Abel HJ, Larson DE, Regier AA, Chiang C, Das I, Kanchi KL et al. Mapping and characterization of structural variation in 17,795 human genomes. Nature 2020; 583: 83–89.

5. Hickey G, Heller D, Monlong J, Sibbesen JA, Sirén J, Eizenga J et al. Genotyping structural variants in pangenome graphs using the vg toolkit. Genome Biol 2020; 21: 35.

6. Sirén J, Monlong J, Chang X, Novak AM, Eizenga JM, Markello C et al. Pangenomics enables genotyping of known structural variants in 5202 diverse genomes. Science 2021; 374: abg8871.

7. Paten B, Eizenga JM, Rosen YM, Novak AM, Garrison E, Hickey G. Superbubbles, Ultrabubbles, and Cacti. J Comput Biol 2018; 25: 649–663.

8. Rautiainen M, Nurk S, Walenz BP, Logsdon GA, Porubsky D, Rhie A et al. Telomere-to-telomere assembly of diploid chromosomes with Verkko. Nat Biotechnol 2023. doi:10.1038/s41587-023-01662-6.

9. Just W. Computational Complexity of Multiple Sequence Alignment with SP-Score. Journal of Computational Biology 2004; 8.https://www.liebertpub.com/doi/10.1089/106652701753307511 (accessed 5 Oct2022).

10. Kille B, Balaji A, Sedlazeck FJ, Nute M, Treangen TJ. Multiple genome alignment in the telomere-to-telomere assembly era. Genome Biol 2022; 23: 1–22.

11. Blanchette M, Kent WJ, Riemer C, Elnitski L, Smit AFA, Roskin KM et al. Aligning multiple genomic sequences with the threaded blockset aligner. Genome Res 2004; 14: 708–715.

12. Harris RS. Improved pairwise alignment of genomic DNA. 2007.https://search.proquest.com/openview/bc77cca0fb9390b44b9ef572fb574322/1?pq-origsite=gscholar&cbl=18750.

13. Armstrong J, Hickey G, Diekhans M, Fiddes IT, Novak AM, Deran A et al. Progressive Cactus is a multiple-genome aligner for the thousand-genome era. Nature 2020; 587: 246–251.

14. Goenka SD, Turakhia Y, Paten B, Horowitz M. SegAlign: A Scalable GPU-Based Whole Genome Aligner. In: SC20: International Conference for High Performance Computing, Networking, Storage and Analysis. IEEE, 2020 doi:10.1109/sc41405.2020.00043.

15. Paten B, Diekhans M, Earl D, St. John J, Ma J, Suh B et al. Cactus Graphs for Genome Comparisons. Journal of Computational Biology 2011; 18: 461–489.

16. Li H, Feng X, Chu C. The design and construction of reference pangenome graphs with minigraph. Genome Biol 2020; 21: 1–19.

17. Lee C, Grasso C, Sharlow MF. Multiple sequence alignment using partial order graphs. Bioinformatics 2002; 18: 452–464.

18. Li H. Minimap2: pairwise alignment for nucleotide sequences. Bioinformatics 2018; 34: 3094–3100.

19. Vivian J, Rao AA, Nothaft FA, Ketchum C, Armstrong J, Novak A et al. Toil enables reproducible, open source, big biomedical data analyses. Nat Biotechnol 2017; 35: 314–316.

20. Paten B, Earl D, Nguyen N, Diekhans M, Zerbino D, Haussler D. Cactus: Algorithms for genome multiple sequence alignment. Genome Res 2011; 21: 1512–1528.

21. Hickey G, Paten B, Earl D, Zerbino D, Haussler D. HAL: a hierarchical format for storing and analyzing multiple genome alignments. Bioinformatics 2013; 29: 1341–1342.

22. Fiddes IT, Armstrong J, Diekhans M, Nachtweide S, Kronenberg ZN, Underwood JG et al. Comparative Annotation Toolkit (CAT)-simultaneous clade and personal genome annotation. Genome Res 2018; 28: 1029–1038.

23. Doerr D. GFAffix. GitHub. 2022.https://github.com/marschall-lab/GFAffix/releases/tag/0.1.3 (accessed 5 Oct2022).

24. Bzikadze AV, Pevzner PA. TandemAligner: a new parameter-free framework for fast sequence alignment. bioRxiv. 2022; : 2022.09.15.507041.

25. Liao W-W, Asri M, Ebler J, Doerr D, Haukness M, Hickey G, et al. A Draft Human Pangenome Reference. bioRxiv. 2022; : 2022.07.09.499321.

26. Nurk S, Koren S, Rhie A, Rautiainen M, Bzikadze AV, Mikheenko A et al. The complete sequence of a human genome. Science 2022; 376: 44–53.

27. Rautiainen M, Marschall T. GraphAligner: rapid and versatile sequence-to-graph alignment. Genome Biol 2020; 21: 253.

28. Poplin R, Chang P-C, Alexander D, Schwartz S, Colthurst T, Ku A et al. A universal SNP and small-indel variant caller using deep neural networks. Nat Biotechnol 2018; 36: 983–987.

29. Wagner J, Olson ND, Harris L, McDaniel J, Cheng H, Fungtammasan A et al. Curated variation benchmarks for challenging medically relevant autosomal genes. Nat Biotechnol 2022; 40: 672–680.

30. Ebler J, Ebert P, Clarke WE, Rausch T, Audano PA, Houwaart T et al. Pangenome-based genome inference allows efficient and accurate genotyping across a wide spectrum of variant classes. Nat Genet 2022; 54: 518–525.

31. 1000 Genomes Project Consortium, Auton A, Brooks LD, Durbin RM, Garrison EP, Kang HM et al. A global reference for human genetic variation. Nature 2015; 526: 68–74.

32. Ebert P, Audano PA, Zhu Q, Rodriguez-Martin B, Porubsky D, Bonder MJ et al. Haplotype-resolved diverse human genomes and integrated analysis of structural variation. Science 2021; 372. doi:10.1126/science.abf7117.

33. Chakraborty M, Emerson JJ, Macdonald SJ, Long AD. Structural variants exhibit widespread allelic heterogeneity and shape variation in complex traits. Nat Commun 2019; 10: 1–11.

34. Huang W, Massouras A, Inoue Y, Peiffer J, Ràmia M, Tarone AM et al. Natural variation in genome architecture among 205 Drosophila melanogaster Genetic Reference Panel lines. Genome Res 2014; 24: 1193–1208.

35. Garrison E, Marth G. Haplotype-based variant detection from short-read sequencing. 2012. doi:10.48550/arXiv.1207.3907.

36. Miller DE, Kahsai L, Buddika K, Dixon MJ, Kim BY, Calvi BR et al. Identification and Characterization of Breakpoints and Mutations on Drosophila melanogaster Balancer Chromosomes. G3 Genes|Genomes|Genetics 2020; 10: 4271–4285.

37. Sherman RM, Forman J, Antonescu V, Puiu D, Daya M, Rafaels N et al. Assembly of a pan-genome from deep sequencing of 910 humans of African descent. Nat Genet 2019; 51: 30–35.

38. hpp_pangenome_resources. Github https://github.com/human-pangenomics/hpp_pangenome_resources (accessed 1 Mar 2023).

39. Guarracino A, Buonaiuto S, Potapova T, Rhie A, Koren S, Rubinstein B, et al. Recombination between heterologous human acrocentric chromosomes. bioRxiv. 2022; : 2022.08.15.504037.

40. Zhou Y, Zhang Z, Bao Z, Li H, Lyu Y, Zan Y et al. Graph pangenome captures missing heritability and empowers tomato breeding. Nature 2022; 606: 527–534.

41. Leonard AS, Crysnanto D, Fang Z-H, Heaton MP, Vander Ley BL, Herrera C et al. Structural variant-based pangenome construction has low sensitivity to variability of haplotype-resolved bovine assemblies. Nat Commun 2022; 13: 1–13.

42. Li H. Identifying centromeric satellites with dna-brnn. Bioinformatics 2019; 35: 4408–4410.

43. Gao Y, Liu Y, Ma Y, Liu B, Wang Y, Xing Y. abPOA: an SIMD-based C library for fast partial order alignment using adaptive band. Bioinformatics 2021; 37: 2209–2211.

44. Garrison E, Guarracino A. Unbiased pangenome graphs. Bioinformatics 2023; 39. doi:10.1093/bioinformatics/btac743.

45. Eizenga JM, Novak AM, Kobayashi E, Villani F, Cisar C, Heumos S et al. Efficient dynamic variation graphs. Bioinformatics 2020; 36: 5139–5144.

46. Sirén J, Garrison E, Novak AM, Paten B, Durbin R. Haplotype-aware graph indexes. Bioinformatics 2020; 36: 400–407.

47. Mose LE, Wilkerson MD, Hayes DN, Perou CM, Parker JS. ABRA: improved coding indel detection via assembly-based realignment. Bioinformatics 2014; 30: 2813–2815.

48. Zook JM, Catoe D, McDaniel J, Vang L, Spies N, Sidow A et al. Extensive sequencing of seven human genomes to characterize benchmark reference materials. Scientific Data 2016; 3: 1–26.

49. Krusche P, Trigg L, Boutros PC, Mason CE, De La Vega FM, Moore BL et al. Best practices for benchmarking germline small-variant calls in human genomes. Nat Biotechnol 2019; 37: 555–560.

50. Li H, Bloom JM, Farjoun Y, Fleharty M, Gauthier L, Neale B et al. A synthetic-diploid benchmark for accurate variant-calling evaluation. Nat Methods 2018; 15: 595–597.

51. broadinstitute/picard. GitHub. https://github.com/broadinstitute/picard (accessed 5 Oct 2022).

52. Kuhn RM, Haussler D, Kent WJ. The UCSC genome browser and associated tools. Brief Bioinform 2012; 14: 144–161.

53. English AC, Menon VK, Gibbs RA, Metcalf GA, Sedlazeck FJ. Truvari: refined structural variant comparison preserves allelic diversity. Genome Biol 2022; 23: 271.

54. Danecek P, Bonfield JK, Liddle J, Marshall J, Ohan V, Pollard MO et al. Twelve years of SAMtools and BCFtools. Gigascience 2021; 10: giab008.

55. Cleary JG, Braithwaite R, Gaastra K, Hilbush BS, Inglis S, Irvine SA, et al. Comparing Variant Call Files for Performance Benchmarking of Next-Generation Sequencing Variant Calling Pipelines. bioRxiv. 2015; : 023754.

56. Smit AFA, Hubley R,, Green, P. RepeatMasker Open-4.0. 2013–2015.http://www.repeatmasker.org.

